# Differential requirement for the EDS1 catalytic triad in *A. thaliana* and *N. benthamiana*

**DOI:** 10.1101/2021.12.15.472806

**Authors:** Josua Zönnchen, Johannes Gantner, Dmitry Lapin, Karen Barthel, Lennart Eschen-Lippold, Stefan Zantop, Carola Kretschmer, Jane E. Parker, Raphael Guerois, Johannes Stuttmann

**Affiliations:** Institute for Biology, Department of Plant Genetics, Martin Luther University Halle-Wittenberg, D-06120 Halle, Germany; Department of Plant-Microbe Interactions, Max-Planck Institute for Plant Breeding Research, D-50829 Cologne, Germany; Plant-Microbe Interactions, Department of Biology, Utrecht University, Utrecht, The Netherlands; Department of Crop Physiology, Institute of Agricultural and Nutritional Sciences, Martin Luther University Halle-Wittenberg, D-06120 Halle,Germany; Department of Biochemistry of Plant Interactions, Leibniz Institute of Plant Biochemistry, D-06120 Halle, Germany; Cologne-Düsseldorf Cluster of Excellence in Plant Sciences (CEPLAS) D-40225 Düsseldorf, Germany; Institute for Integrative Biology of the Cell (I2BC), IBITECS, CEA, CNRS, Université Paris-Saclay, F-91198, Gif-sur-Yvette, France; Institute for Biosafety in Plant Biotechnology, Federal Research Centre for Cultivated Plants, Julius Kühn-Institute (JKI), Quedlinburg, Germany

## Abstract

- Heterodimeric complexes incorporating the lipase-like proteins EDS1 with PAD4 or SAG101 are central hubs in plant innate immunity. EDS1 functions encompass signal relay from TIR domain-containing intracellular NLR-type immune receptors (TNLs) towards RPW8-type helper NLRs (RNLs) and, in *A. thaliana*, bolstering of signaling and resistance mediated by cell-surface pattern recognition receptors (PRRs). Biochemical activities underlying these mechanistic frameworks remain unknown.
- We used CRISPR/Cas-generated mutant lines and agroinfiltration-based complementation assays to interrogate functions of EDS1 complexes in *N. benthamiana*.
- We do not detect impaired PRR signaling in *N. benthamiana* lines deficient in EDS1 complexes or RNLs. Intriguingly, mutations within the catalytic triad of *Solanaceae* EDS1 can abolish or enhance TNL immunity in *N. benthamiana*. Furthermore, nuclear EDS1 accumulation is sufficient for *N. benthamiana* TNL (Roq1) immunity.
- Reinforcing PRR signaling in Arabidopsis might be a derived function of the TNL/EDS1 immune sector. Dependency of *Solanaceae* but not *A. thaliana* EDS1 on catalytic triad residues raises the possibility that a TNL-derived small molecule binds to the *Solanaceae* EDS1 lipase-like domain, and that EDS1 lipase-like domain pocket contributions to TNL immune responses vary between lineages. Whether and how nuclear EDS1 activity connects to membrane pore-forming RNLs remains unknown.

## Introduction

Genetically encoded cell surface-resident and intracellular immune receptors serve as detection devices in plant and animal innate immunity. In plants, which lack an adaptive immune system, immune receptor repertoires have expanded and diversified between different species or even accessions (Barragan and Weigel, 2021). At the cell surface, pattern recognition receptors (PRRs) detect microbe-associated molecular patterns (MAMPs) to induce pattern-triggered immunity (PTI). Inside the cell, receptors of the nucleotide-binding/leucine rich repeat (NLR) class detect effector proteins (virulence factors secreted into the host cell cytoplasm by pathogenic microbes) to induce effector-triggered immunity (ETI).

PRRs are receptor-like kinases (RLKs) or receptor-like proteins (RLPs) with diverse ectodomains (reviewed in Saijo et al., 2018). RLKs and RLPs associate, either constitutively or in a stimulus-dependent manner, with co-receptors, such as the RLKs BAK1/SERK3 (Brassinosteroid-insensitive 1 (BRI1) Associated Kinase1/Somatic Embryogenesis Receptor Kinase 3) or SOBIR1 (Suppressor Of BIR1,1; Chinchilla et al., 2007; Heese et al., 2007; Liebrand et al., 2013). PRR-ligand binding induces a suite of downstream signaling events and physiological responses, including activation of mitogen-activated protein kinases (MAPKs), Ca^2+^-influx, production of reactive oxygen species (ROS burst) and ethylene, and induction of defense genes (Saijo et al., 2018). Some PRRs detect pathogen isolate (strain)-specific ligands, such as apoplastic effectors of the fungal pathogens *Verticillium dahliae* and *Cladosporium fulvum*, and induce strong resistance responses accompanied by programmed cell death (Thomma et al., 2011). However, most characterized MAMPs are relatively conserved and widely distributed molecules characteristic for a class or group of organisms, such as fungal chitin or peptides derived from bacterial flagellin (Saijo et al., 2018). Activation of the respective PRRs is normally not accompanied by cell death. Accordingly, PTI is generally considered a low level resistance response sufficient to combat non-adapted pathogens.

Host-adapted pathogens that overcome PTI confront the ETI defense layer. A rapid and strong ETI response, which is often accompanied by programmed cell death at infection sites (the hypersensitive response, HR), is induced in the presence of an effector and a cognate NLR-type immune receptor. The canonical NLR architecture consists of a C-terminal leucine-rich repeat (LRR), a central nucleotide-binding (NB)/oligomerization and an N-terminal signaling domain (Bentham et al., 2017). Plant NLRs are subdivided into three major classes based on their N-terminal domains: TNLs carrying a Toll-interleukin 1 receptor (TIR) domain, CNLs a coiled coil (CC) domain, and RNLs a CC_R_ or HeLo domain, a subtype of the CC domain also found in the non-NLR immunity regulators RPW8 and MLKL (Resistance to Powdery Mildew 8 and Mixed Lineage Kinase-Like; Xiao et al., 2005; Collier et al., 2011; Lapin et al., 2020; Mahdi et al., 2020). Most characterized CNLs and TNLs function as sensor NLRs (sNLRs) in pathogen effector detection. Plant sNLR repertoires are diverse and can range from a few to several hundred *NLR* genes (Baggs et al., 2020). In contrast, RNLs are more conserved and operate in basal resistance against virulent pathogens and as helper NLRs (hNLRs) downstream of TNLs and some CNLs (Bonardi et al., 2011; Castel et al., 2018; Wu et al., 2018; Lapin et al., 2019; Saile et al., 2020; Sun et al., 2021). In agreement with broader and potentially ancestral immune functions, two conserved subgroups of RNLs, ADR1 (Activated Disease Resistance 1) and NRG1 (N-Required Gene 1) RNLs, were detected in the genomes of nearly all flowering plants (Collier et al., 2011).

While ADR1s are present in seed plant genomes, NRG1s are restricted to eudicots with expanded TNL panels (Collier et al., 2011; Lapin et al., 2020; Liu et al., 2021). Recent reports suggest that sensor CNLs and RNL-type hNLRs form oligomers (resistosomes) upon activation, which can insert into membranes and function as cation-permeable channels (Wang et al., 2019; Bi et al., 2021; Jacob et al., 2021). Although it remains unclear whether resistosome formation by *Arabidopsis thaliana* (Arabidopsis) CNL ZAR1 (HopZ-Activated Resistance 1) is prototypical for CNLs and RNLs, it is possible that Ca2^+^ influx represents a common output in CNL and TNL-RNL immunity.

PTI and ETI were traditionally considered as independent immune sectors contributing to pathogen resistance and converging on transcriptional defenses (Tao et al., 2003; Cui et al., 2015). Recent reports suggest that ETI and PTI cross-potentiate each other in pathogen resistance (Lu and Tsuda, 2021). PTI-deficient Arabidopsis lines failed to mount efficient ETI responses (Ngou et al., 2021; Yuan et al., 2021). Reciprocally, ETI components were required for PTI in Arabidopsis: Lines deficient in a central regulator of TNL immunity, EDS1 (Enhanced Disease Susceptibility 1) or RNLs were unable to mount full TNL immunity and were impaired in early and late PTI responses (Pruitt et al., 2021; Tian et al., 2021). Evidence of PTI-ETI connectivity is so far mainly limited to Arabidopsis and the underlying mechanisms remain unknown. For example, it is unclear whether EDS1 contributes directly to PTI signaling or whether PTI signaling is primed and reinforced by TNL signaling *via* EDS1 (Pruitt et al., 2021; Tian et al., 2021).

EDS1 is part of a small plant-specific protein family containing also PAD4 (Phytoalexin-Deficient 4) and SAG101 (Senescence Associated Gene 101; Lapin et al., 2020). EDS1 family proteins are characterized by the fusion of an α/ß-hydrolase (class-3 lipase) domain with a C-terminal EP (EDS1-PAD4) all-helical domain which has no significant similarities to any known structure (Wagner et al., 2013; Bhandari et al., 2019). EDS1 forms mutually exclusive heterodimers with PAD4 or SAG101 (Wagner et al., 2013). In Arabidopsis, the EDS1-PAD4 complex has a major role in TNL-mediated ETI and reinforces signaling by surface receptors (Dongus and Parker, 2021). By contrast, EDS1-SAG101 dimers promote TNL-triggered defense and host cell death (Feys et al., 2005; Wagner et al., 2013; Lapin et al., 2019). Distinct Arabidopsis EDS1-PAD4-ADR1 and EDS1-SAG101-NRG1 modules regulating pathogen resistance and cell death, respectively, were recently proposed based on genetic and biochemical analyses (Lapin et al., 2019; Sun et al., 2021; Wu et al., 2021).

In our current understanding, EDS1 complexes operate downstream of TNL receptors but upstream of RNLs, since autoactive RNL fragments, but not TNL activation or autoactive TNLs and isolated TIR domains, can induce cell death and resistance signaling in *eds1* mutant plants (Qi et al., 2018; Horsefield et al., 2019; Wan et al., 2019; Jacob et al., 2021). Hence, EDS1 complexes probably relay a signal from activated TNLs to RNLs. Upon activation, the TNLs RPP1 (Recognition Of *Peronospora Parasitica* 1) and Roq1 (Recognition Of XopQ 1) tetramerize into holoenzymes with NADase activity (Ma et al., 2020; Martin et al., 2020). This suggests that signal relay is mediated by a small molecule, although the *in planta* TNL NADase active products remain unknown (Wan et al., 2019; Duxbury et al., 2020; Yu et al., 2021). This hypothesis appears even more plausible because both EDS1 dimers form an EP-domain cavity lined by conserved residues that are essential for signaling (Bhandari et al., 2019; Gantner et al., 2019; Lapin et al., 2019; Sun et al., 2021). Moreover, EDS1 and PAD4 (but not SAG101) orthologs have conserved N-terminal lipase-like domain pockets with a characteristic serine-aspartate-histidine catalytic triad which could bind and/or process a TNL-generated small molecule (Wagner et al., 2013; Voss et al., 2019). Notably, mutational analyses revealed that an intact lipase-domain enzymatic pocket of Arabidopsis EDS1-PAD4 complexes is dispensable for TNL immunity but necessary for PAD4-mediated resistance to aphid attack (Louis et al., 2012; Wagner et al., 2013; Dongus et al., 2020).

We established *Nicotiana benthamiana* (*Nb*) as a genetic system for analysis of EDS1-family functions in TNL immunity (Adlung et al., 2016; Ordon et al., 2017; Gantner et al., 2019). Our analyses revealed that, in contrast to Arabidopsis, an EDS1-SAG101b complex (most *Solanaceae* genomes encode two SAG101 isoforms) is necessary and sufficient for all tested TNL-mediated immune responses in *Nb*; immune functions of *Nb*PAD4 were not detected (Gantner et al., 2019). In agreement with an EDS1-SAG101-NRG1 module conferring TNL ETI in *Nb* (Lapin et al., 2019; Sun et al., 2021), a TNL immune response was also largely abolished in NRG1-deficient *Nb* plants (Qi et al., 2018), arguing against a separation of pathogen resistance and cell death at the level of EDS1-RNL complexes in this system.

In this study, we investigated EDS1 dimer functions and the subcellular compartments in which EDS1 complexes are localized during *Nb* PTI and ETI responses. Our data suggest that recruitment to PTI signaling does not represent a general function of EDS1-RNL modules. In *Nb* but not Arabidopsis, amino acid exchanges within the EDS1 catalytic triad can enhance or suppress EDS1-dependent TNL immune responses, but the critical enzymatic serine residue is not required. These data are compatible with binding of a small molecule during EDS1 signaling, although further mutational analyses do not support small molecule binding in proximity to the presumed EDS1 active site. Also, our data suggest that mainly nuclear EDS1 complexes mediate immune signaling during an *Nb* ETI response. This puts into question the compartment in which RNLs, that are proposed to form plasma membrane pores, are activated and signal in *Solanaceae* TNL immunity.

## Results

### EDS1 complexes are dispensable for signaling by tested cell surface receptors in *N. benthamiana*

The EDS1-PAD4-ADR1 module fulfills a major immune function in Arabidopsis TNL-mediated pathogen resistance and contributes to PRR signaling (Wagner et al., 2013; Lapin et al., 2019; Pruitt et al., 2021; Tian et al., 2021). Although *NbPAD4* gene expression was induced upon pathogen challenge and *Sl*EDS1-*Sl*PAD4 from tomato (*Solanum lycopersicum*, *Sl*) can mediate TNL immunity upon transfer to Arabidopsis, *Nb pad4* mutant plants were not impaired in TNL ETI assays (Gantner et al., 2019; Lapin et al., 2019). We therefore investigated whether the EDS1-PAD4 module or EDS1 complexes together with RNLs contribute to PRR signaling in *Solanaceae*, as previously suggested (Hu et al., 2005; Gabriels et al., 2007).

We first generated an *Nb bak1*/*serk3* mutant line (referred to as *Nb bak1*) as a negative control for PRR signaling. In the *Nb bak1* line, two LRR-RLK-coding genes previously silenced by Heese et al. (Heese et al., 2007; Wang et al., 2018) were disrupted by genome editing (Fig S1, Table S1). *Nb bak1* mutant plants developed cell death similar to wild type upon expression of several different effectors or a TIR fragment of the Arabidopsis TNL DM2h (Dangerous Mix 2h; Ordon et al., 2021), suggesting that cell death pathways triggered intracellularly are not impaired (Fig S2).

Alongside wild type and *Nb bak1* plants, we tested induction of host cell death after activation of the tomato LRR-RLPs Cf-4 and Cf-9 (*Cladosporium fulvum*-4/-9) in *Nb* mutant lines deficient in EDS1 complexes (*eds1* or *pad4 sag101a sag101b* (*pss*) triple mutant) or the RNL NRG1. Cf4 and Cf9 can be activated by transient co-expression (*via* agroinfiltration) of their respective *C. fulvum* ligands, Avr4 and Avr9 (Avirulence 4/9) in *Nb* (Van der Hoorn et al., 2000). Co-expression of the receptor-ligand pairs, but not either component alone, induced cell death in wild-type *Nb*, as expected (Figs 1a, S3a). Using low inoculum densities (see figure legends), we observed reduced cell death on *Nb bak1* but not mutant lines deficient in EDS1 complexes or the RNL NRG1 (Figs 1a, S3a). We quantified cell death by ion leakage assays (Figs 1b, S3b). *Nb bak1* but none of the EDS1 complex or RNL-deficient lines displayed lower ion leakage in the Cf4/Avr4-induced response (Fig 1b). Cf9/Avr9-induced cell death is generally weaker (Van der Hoorn et al., 2000) and we did not detect significant differences in ion leakage between lines, which was overall low and fluctuating (Fig S3b).

**Figure 1:**
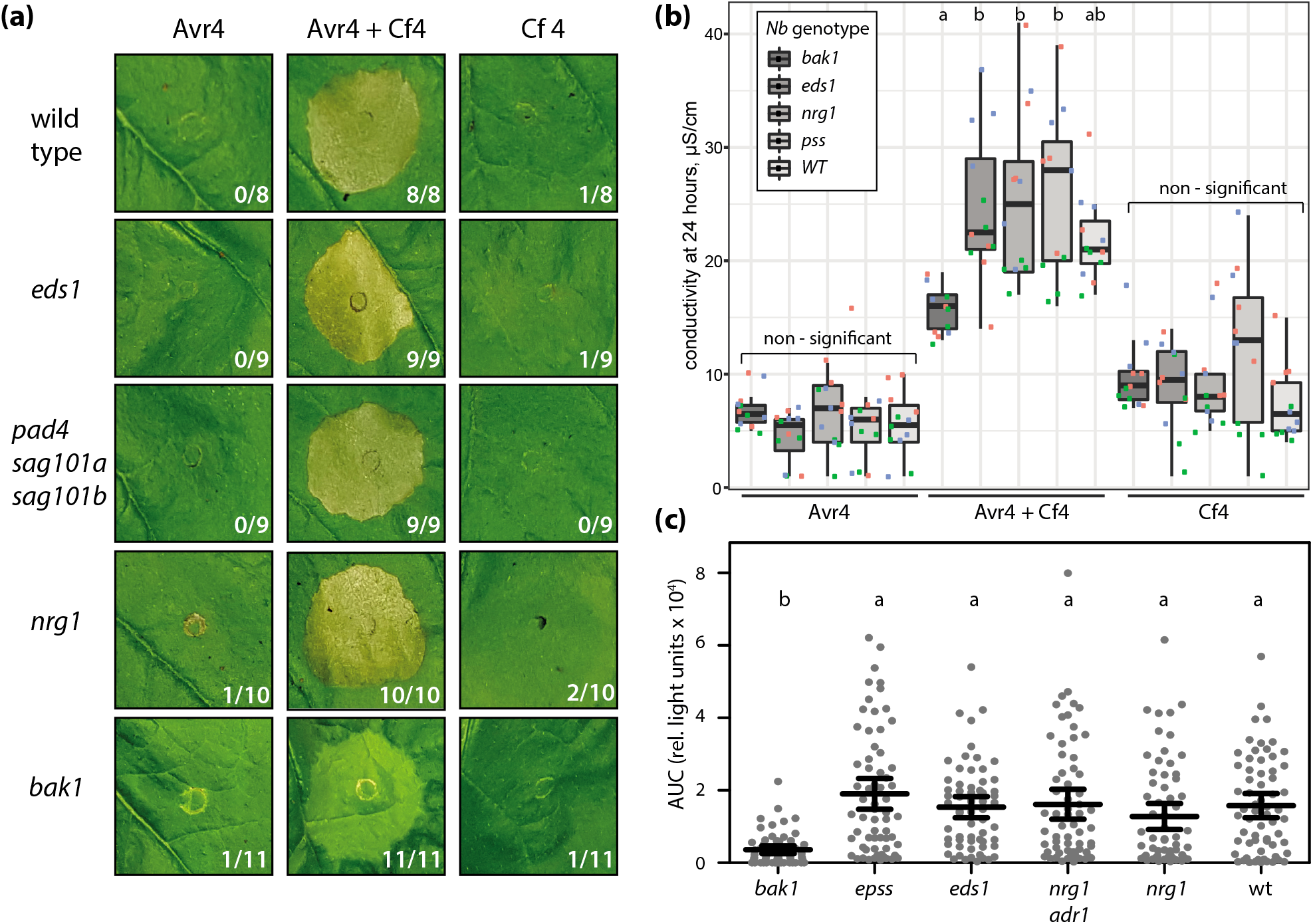
RLP- and RLK-mediated responses in *N. benthamiana* lines lacking EDS1 complexes or RNLs. **(a)** Cell death induction upon (co-)expression of Avr4 and Cf-4. Avr4 and/or Cf-4 were expressed by agroinfiltration (OD_600_ = 0.05) in the indicated *Nb* lines. Symptom (cell death) formation was documented 4 dpi. The experiment was conducted 8 times with similar results, and representative images are shown. Numbers indicate infiltration sites with chlorosis (*bak1*) or cell death (all remaining genotypes). **(b)** Quantitative assessment of cell death by ion leakage measurements. Leaf discs were sampled 24 hpi, washed for 2h in H_2_O, and incubated 24h in H_2_O under shaking prior to measuring conductivity. Three independent experiments, each conducted with four replicates, are shown. Letters indicate statistically significant differences (ANOVA, Tukey HSD, p < 0.001). **(c)** flg22-induced ROS production in different *Nb* lines. Leaf discs of 4 week-old *Nb* plants of the indicated genotypes were treated with 2 nM flg22, and ROS production was measured over 60 minutes. The area under the curve for 64 measurements from four independent experiments was plotted (mean, 95% confidence interval). Letters indicate statistically significant differences (ANOVA, Bonferroni *post-hoc* test, p < 0.05).

We next tested PTI responses initiated by LRR-RLK *Nb*FLS2 in the *Nb* mutant lines, including a line lacking all EDS1 proteins (*epss* quadruple mutant; Lapin et al., 2019) and a newly generated *Nb nrg1 adr1* double mutant (Fig 1c; Prautsch et al., 2021). By dilution series, we determined 2 nM flg22 as the minimal elicitor concentration for inducing a reliable ROS burst in wild-type plants under our conditions. In corresponding assays of the mutant lines, flg22-elicited ROS production was significantly reduced in *Nb bak1*, but not in any other mutant (Fig 1c).

In summary, we detected reduced cell death (Cf4/9) and ROS burst (*Nb*FLS2) in *Nb bak1*, but not a contribution of PAD4, EDS1 complexes or RNLs to the tested LRR-RLPs Cf-4 and Cf-9 and LRR-RLK FLS2. The data show that neither EDS1 dimers nor RNLs are essential for the tested cell surface receptor-triggered immune responses in *Nb*. We concluded that recruitment of these components for PRR signaling probably does not represent a conserved function of EDS1, TNLs or RNLs between Arabidopsis and *Nb*.

### The EDS1 catalytic triad is critical for Roq1 TNL immunity in *N. benthamiana*

The EDS1-SAG101b complex mediates all known TNL immune functions in *Nb* and the C-terminal EP domains of both proteins are essential for TNL signaling (Gantner et al., 2019; Lapin et al., 2019). Here, we interrogated functions of the *Sl*EDS1 N-terminal lipase-like domain.

In the functional *Sl*EDS1-*Sl*SAG101b complex, *Sl*EDS1 but not *Sl*SAG101b, contains a conserved S-D-H catalytic triad (Fig 2a; Wagner et al., 2013; Gantner et al., 2019). The serine residue (S125 in *Sl*EDS1) is embedded in a GXSXG motif forming the so-called nucleophile elbow (Fig 2a). The serine nucleophile is involved in formation of the first reaction intermediate in a canonical α/ß-hydrolase mechanism, which also requires an oxyanion hole (Rauwerdink and Kazlauskas, 2015). Previous analyses of *At*EDS1 (in complex with *At*SAG101) revealed high spatial conservation of the triad residues, whereas the oxyanion hole was disturbed in the crystal structure (Wagner et al., 2013).

**Figure 2:**
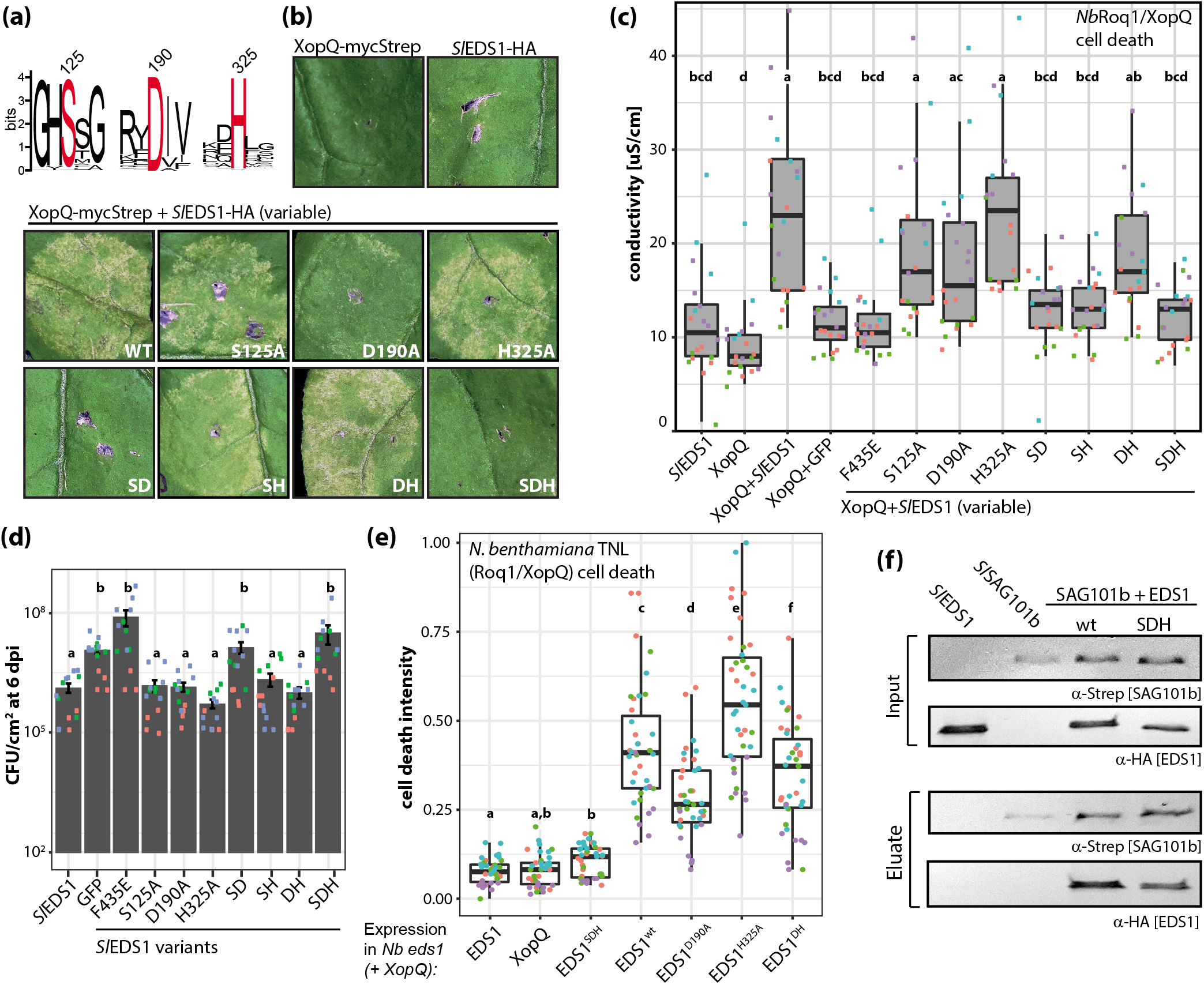
The *Sl*EDS1 catalytic triad is critical for XopQ-induced cell death and pathogen resistance in *N. benthamiana*. **(a)** Sequence logo of the predicted EDS1 catalytic triad positions generated from an alignment of 75 orthologous sequences. The triad S-D-H residues are highlighted, numbering corresponds to *Sl*EDS1. EDS1 sequences from Lapin et al. (2019). **(b)** Cell death induction upon (co-)expression of XopQ and *Sl*EDS1-variants as indicated. Agrobacterium strains for the expression of indicated proteins (under 35S promoter control) were infiltrated at OD_600_ = 0.4 per strain into *Nb eds1* mutant plants. At 3 dpi, samples were taken for verification of protein expression (Fig S4), and symptom (cell death) formation was documented 7 dpi. The experiment was conducted >10 times with similar results, and representative images are shown. **(c)** Quantification of cell death by ion leakage measurements. Infiltrations and strains as in (a), but at 4 dpi, samples were taken for ion leakage measurements. The experiment was conducted four times with 4-6 replicates; independent experiments are color-coded. Letters indicate statistically significant differences (ANOVA, Tukey HSD, p<0.05). **(d)** *Xcv* growth assay in *Nb eds1* plants transiently expressing GFP (negative control), *Sl*EDS1 (positive control) or variants thereof, as indicated. Plants were co-infiltrated with Agrobacterium strains and *Xcv* bacteria, and *Xcv* bacterial titers determined 6 dpi. The experiment was conducted three times with 4-6 replicates. Individual data points are color-coded for experiments. Error bars represent SEM. Letters indicate statistically significant differences (ANOVA, Tukey HSD, p<0.05). **(e)** Quantification of cell death by red light imaging. Graph shows composite data originating from four independent experiments. Per experiment, 7 - 12 leaves were used for agroinfiltration and documented 5 dpi. HR cell death was quantified using ImageJ and normalized to a 0-1 range. Letters indicate statistically significant differences (ANOVA, Fisher LSD, p < 0.05). **(f)** Co-purification assay with *Sl*SAG101b-mycStrepII and *Sl*EDS1-6xHA or variants thereof. Indicated proteins were, separately or simultaneously, expressed in *Nb*. Samples were taken 3 dpi, and subjected to Strep purification. Input and eluate fractions were analyzed by SDS-PAGE and immunodetection. The experiment was conducted four times with similar results. A co-purification assay including single amino acid exchange variants of *Sl*EDS1 is shown in Figure S4b.

In *Nb*, TNL immunity and cell death can be induced by Agrobacterium-mediated transient expression of the *Xanthomonas campestris* pv. *vesicatoria* (*Xcv*) effector XopQ (*Xanthomonas* Outer Protein Q), recognized by TNL Roq1 (Schultink et al., 2017). In XopQ cell death assays, mutant lines deficient in *EDS1* family genes can be transiently complemented by co-expression of respective proteins from tomato (*Sl*) or *Nb* with XopQ (Gantner et al., 2019; Lapin et al., 2019). Also, EDS1-dependent pathogen resistance can be measured by mixed Agrobacterium-*Xcv* infections (Lapin et al., 2019).

We generated *Sl*EDS1 variants with single or combined exchanges (to alanine) of the catalytic triad residues S125, D190 and H325. The *Sl*EDS1 variants were first tested in cell death assays by co-expression with XopQ in *Nb eds1* mutant plants (Fig 2b). We consistently observed reduced TNL-triggered cell death upon co-expression of EDS1^D190A^ with XopQ and cell death was abolished with EDS1^SD^ or EDS1^SDH^ (Fig 2b). Exchange of the single S125 residue did not impair *Sl*EDS1 function. Remarkably, co-expression of H325 variants with XopQ induced stronger cell death (Fig 2b; *Sl*EDS1 *vs. Sl*EDS1^H325A^ and *Sl*EDS1^D190A^ *vs*. *Sl*EDS1^DH^). Enhanced or abolished cell death did not correlate to differences in protein accumulation (Fig S4).

Quantification of cell death in ion leakage assays and testing of *Sl*EDS1 variants in *Xcv* resistance assays led to similar results: *Sl*EDS1^SD^ and *Sl*EDS1^SDH^ variants lost all immune activity (Fig 2c,d), similar to the non-functional *Sl*EDS1^F435E^ variant (Gantner et al., 2019). Minor macroscopic effects of *Sl*EDS1^D190A^ and *Sl*EDS1^H325A^ exchanges were not detected in ion leakage assays (Fig 2c,d). Therefore, we quantified the intensity of cell death responses using red-light imaging (Landeo Villanueva et al., 2021). Composite data from multiple (>35) infiltrated leaves confirmed reduced cell death intensities for *Sl*EDS1^D190A^ and more intense cell death with *Sl*EDS1^H325A^ within *Sl*EDS1^H325A^ and *Sl*EDS1^DH^ compared to controls (Fig 2e). The tested *Sl*EDS1 mutant proteins accumulated to similar levels in cell death assays (Fig S4a). All *Sl*EDS1 variants formed a dimer with *Sl*SAG101, tested by co-purification with Strep-tagged *Sl*SAG101b (Figs 2f, S4b). In these experiments, reduced levels of *Sl*EDS1^SDH^ co-purified with Strep-tagged *Sl*SAG101b in most replicates (Fig 2f). Levels of co-purified EDS1^SDH^ exceeded those of *Sl*EDS1^LLI^ (Fig S4b), a variant carrying mutations in the hydrophobic αH helix required for EDS1 stable interaction with SAG101 and functional in XopQ cell death assays (Gantner et al., 2019). Thus, disruption of EDS1-SAG101 complexes or reduced stability of *Sl*EDS1 triad variants does not explain their immunity defects. The results show that mutations within the *Sl*EDS1 catalytic triad can lead to reduced or enhanced immunity function. Put together with our finding that an *Sl*EDS1^S125A^ exchange does not impair TNL ETI, the data suggest that *Sl*EDS1 immune functions are more likely to involve binding than hydrolysis of a putative small molecule to modulate *Sl*EDS1-*Sl*SAG101b dimer function.

### The catalytic triad of Arabidopsis *At*EDS1 is dispensable for immune signaling in Arabidopsis and *N. benthamiana*

Earlier mutagenesis studies in the Arabidopsis system did not detect a contribution of the catalytic triad of EDS1 proteins to TNL signaling (Wagner et al., 2013). In particular, resistance to *Hyaloperonospora arabidopsidis* isolate Cala2 (*Hpa* Cala2) mediated by RPP2 TNLs (Sinapidou et al., 2004) was fully restored in transgenics expressing *At*EDS1^SDH^-*At*PAD4^S118A^ in the *eds1-2 pad4-1* background. Also, an *At*EDS1^SDHFV^ variant, in which the triad environment was further perturbed by F47W and V189M mutations, was functional when expressed in Col *eds1-2* (Wagner et al., 2013).

The contrasting results obtained for *Sl*EDS1 in TNL Roq1 signaling (Fig 2) prompted us to test resistance functions of *At*EDS1 and *At*PAD4 triad mutants in more detail. We used three different approaches (Fig 3a): i) stable transformation of an Arabidopsis *eds1-12 pad4-1 sag101-3* triple mutant (*At eps*; to avoid confounding effects of *SAG101*) with constructs encoding *At*EDS1 and *At*PAD4 or variants, and challenge of T_1_ plants with *Hpa* Cala2; ii) *Xcv*-Roq1 resistance assays in *Nb epss* plants transiently expressing *At*NRG1.1, *At*SAG101 and *At*EDS1 or variants by co-infiltration of plants with Agrobacterium strains and *Xcv* bacteria (Lapin et al., 2019); and iii) Roq1 cell death assays by expressing XopQ with the same set of Arabidopsis proteins in *Nb epss* plants, and ion leakage measurements (Lapin et al., 2019).

**Figure 3:**
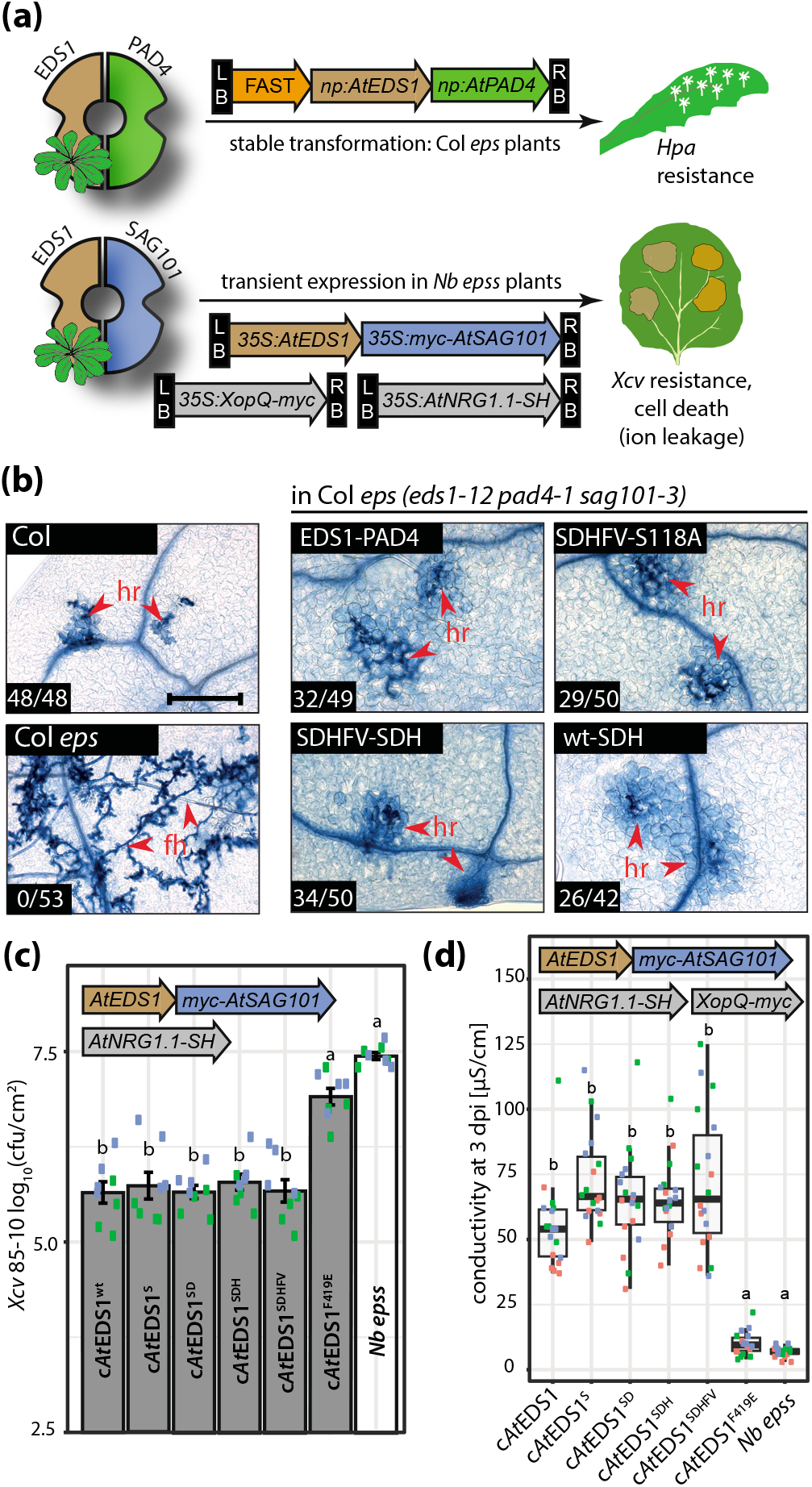
Immune signaling by *At*EDS1-*At*PAD4 complexes does not depend on integrity of the catalytic triad. **(a)** Schematic depiction of used experimental setups to test functionality of Arabidopsis EDS1 family proteins in Arabidopsis (top) and *Nb* (bottom). Constructs coding for Arabidopsis EDS1-PAD4 and variants thereof, under native promoter control, were transformed into Arabidopsis plants deficient in EDS1 family proteins (Col *eps* = *eds1-12 pad4-1 sag101-3*). Resulting primary transformants were directly tested for resistance to *Hpa* isolate Cala2 (panel b). Constructs coding for Arabidopsis EDS1-SAG101 under 35S promoter control were used for transient expression in *N. benthamiana epss* mutant plants. Capacity of variants to mediate pathogen resistance was tested by mixed Agrobacterium-*Xcv* infiltrations with Agrobacterium strains harboring plasmids for expression of *At*EDS1 (or variants thereof), *At*SAG101 and *At*NRG1.1 (panel c). Capacity of variants to mediate cell death was tested by co-expression with *At*SAG101, *At*NRG1 and XopQ and ion leakage measurements (panel d). **(b)** Infection of primary (T_1_) transformants and control lines with *Hpa* isolate Cala2. Primary transformants were selected by seed fluorescence. Three week-old plants were *Hpa*-infected, and first true leaves were used for Trypan Blue staining 6 dpi. Numbers indicate the fraction of plants macroscopically scored as„resistant” and the total number of analyzed plants. At least six primary transformants per construct were analyzed by Trypan Blue staining, and representative micrographs are shown. The experiment was conducted twice with similar results. Scale bar = 250 μm. hr = hypersensitive response; fh = free hyphae. EDS1-SDHFV: S123A/D187A/H317A/F47W/V189M. PAD4-SDH: S118A/D178A/H229A. **(c)** Functionality of *At*EDS1 variants in *Xcv* resistance assays. Bacterial titers were determined 6 dpi. The experiment was conducted twice with 4 replicates in each experiment. Data points from individual experiments are color-coded. Error bars indicate SEM, letters statistically significant differences (ANOVA, Tukey HSD, p < 0.001). **(d)** Functionality of *At*EDS1 variants in cell death induction in *Nb*. Ion leakage was determined 3 dpi as quantitative measurement of cell death. The experiment was conducted three times with six replicates. Error bars and statistics as in **(c)**.

For Arabidopsis *Hpa* Cala2 resistance assays, *At eps* plants were transformed with a single T-DNA for expression of *At*EDS1 with *At*PAD4 under own promoter control. Different combinations of wild-type *At*EDS1, *At*EDS1^SDHFV^ with *At*PAD4, *At*PAD4^S118A^ and *At*PAD4^SDH^ (S118A/D178A/H229A) were tested. In T_1_ resistance assays, plants expressing *At*EDS1^SDHFV^ and *At*PAD4^SDH^ triad variants were as resistant to *Hpa* Cala2 as plants expressing wild-type *At*EDS1-*At*PAD4, and were similar to Col-0 plants (Fig 3b). Thus, all *At*EDS1 and *At*PAD4 lipase catalytic triad variants fully restored RPP2 resistance in the *At eps* mutant background, consistent with earlier data (Wagner et al., 2013).

We obtained similar results in *Xcv*-Roq1 resistance and XopQ-Roq1 cell death assays which recorded immune activities of the *At*EDS1-*At*SAG101 complex in *Nb epss* plants. In both assays, resistance responses mediated by *At*EDS1^SDHFV^ and other triad variants were similar to those of wild-type *At*EDS1 (Fig 3c,d). The non-functional *At*EDS1^F419E^ variant failed to confer resistance or cell death in these experiments (Fig 3c,d). These data argue against a role of the *At*EDS1-*At*PAD4 catalytic triad environments when used to reconstitute TNL immunity in *Nb*.

### Catalytic triads of tomato *Sl*EDS1-*Sl*PAD4 are not required for Arabidopsis TNL (RPS4-RRS1) immunity

We anticipated that the requirement for an intact catalytic triad might be specific for *Sl*EDS1, while *At*EDS1 might function by a different mechanism in Arabidopsis and *Nb*. Therefore, we capitalized on the functionality of the matching *Sl*EDS1-*Sl*PAD4 dimer in Arabidopsis TNL resistance (Gantner et al., 2019; Lapin et al., 2019). *At eps* plants were transformed with constructs co-expressing *Sl*EDS1-GFP with *Sl*PAD4-6xHA or respective triad variants, under control of Arabidopsis promoter fragments (Fig 4a). Functionality of the *Sl*EDS1-*Sl*PAD4 variants was assessed in T_2_ generation transgenic plants using *Pseudomonas syringae pv. tomato* DC3000 (*Pst*) *AvrRps4* that triggers the TNL receptor pair RPS4-RRS1 (Resistant to *P. Syringae* 4 - Resistant to *Ralstonia Solanacearum* 1) (Narusaka et al., 2009; Saucet et al., 2015). We tested 3 to 4 independent transgenic lines for *At eps* expressing *Sl*EDS1-*Sl*PAD4, *Sl*EDS1^SDH^-*Sl*PAD4, *Sl*EDS1-*Sl*PAD4^SDH^, or *Sl*EDS1^SDH^-*Sl*PAD4^SDH^. Plants expressing the wild type *Sl*EDS1-*Sl*PAD4 construct almost fully restored RRS1-RPS4 resistance in the susceptible *At eps* mutant, as expected (Fig 4b; Lapin et al., 2019). All mutant variant constructs restored TNL resistance to the same extent as wild type *Sl*EDS1-*Sl*PAD4 (Fig 4b). Immunodetection assays indicated that the different *Sl*EDS1-*Sl*PAD4 variants were expressed in the transgenic lines (Fig 4c). These experiments show that an intact *Sl*EDS1 catalytic triad is dispensable for TNL RRS1-RPS4 resistance signaling in Arabidopsis, in contrast to its measurable contribution to TNL *Roq1* resistance in *Nb*.

**Figure 4:**
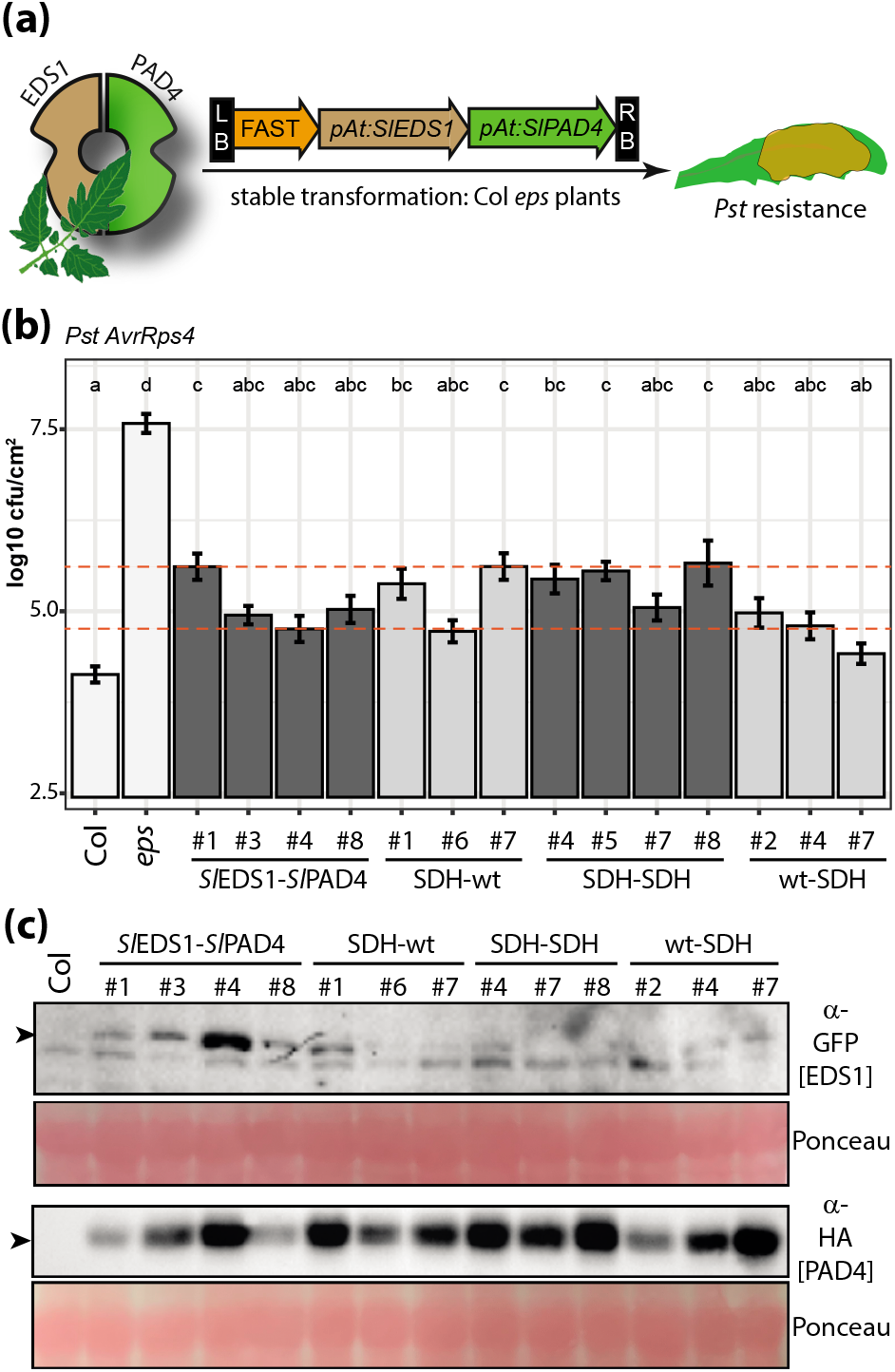
Integrity of the *Sl*EDS1-*Sl*PAD4 catalytic triad is not critical for Arabidopsis TNL immunity. **(a)** Schematic depiction of experimental setup. Constructs coding for tomato EDS1-PAD4 and variants thereof, under control of the respective Arabidopsis promoter elements, were transformed into Arabidopsis plants deficient in EDS1 family proteins (Col *eps* = *eds1-12 pad4-1 sag101-3*). Resulting transformants were tested for resistance to *Pst AvrRps4* bacteria. **(b)** Resistance to *Pst AvrRps4* of transgenic lines expressing *Sl*EDS1-*Sl*PAD4 or triad mutant variants thereof under control of Arabidopsis promoter elements in the Col *eps* mutant background. Transgenic segregants were selected by seed fluorescence (FAST marker) from T_2_ populations. Four week-old plants were syringe-infiltrated with *Pst* AvrRps4, and bacterial titers were determined 3 dpi. At least three independent transgenic lines were tested per transformation, as indicated. Error bars represent SEM; graph shows composite data of 3 independent replicates (18 data points). Letters indicate statistically significant differences (ANOVA, Tukey HSD, p<0.05). **(c)** Immunodetection of *Sl*EDS1-*Sl*PAD4 in Arabidopsis transgenics. Transgenic lines as in b) except SDH-SDH line #5, which was excluded. Five week-old, unchallenged plants were used for immunodetection.

### Perturbation of the *Sl*EDS1 catalytic triad environment or lid region does not impair immune functions

We wanted to challenge further the hypothesis that the *Sl*EDS1 lipase-like domain catalytic pocket binds a small molecule in the *Nb* system. EDS1 possesses an extended lid domain, composed of the helices αF, αG and αH (Wagner et al., 2013). αG and αH are in direct contact with PAD4 or SAG101 in complexes, whereas αF lies across the entrance to a putative catalytic pocket (Figs 5a,S5a). The substrate binding site of lid-containing α/β-hydrolases is commonly composed of residues belonging to the core α/β-hydrolase and the lid domain, as observed in rice gibberellic acid (GA) receptor GID1 (Gibberellin Insensitive Dwarf 1; Shimada et al., 2008; Rauwerdink and Kazlauskas, 2015). To perturb binding of a potential substrate to *Sl*EDS1, we first targeted amino acids in vicinity of the *Sl*EDS1 catalytic triad for mutagenesis. For this we generated *Sl*EDS1 I214E, F235C, F235S, R194A, R194L, W58S, V192A, M195A, M195E, I210A, and I214T variants (Figs S5, 5a). *Sl*EDS1 variants were co-expressed together with XopQ in XopQ/Roq1 cell death assays in *Nb eds1* plants (Fig 5a). Surprisingly, all variants mediated Xoq/Roq1 cell death as efficiently as wild-type *Sl*EDS1. We next generated *Sl*EDS1 variants in which residues comprising predicted helix αF were deleted, and joined to αG by short linker sequences. In lid variants lidΔ#1 and lidΔ#2, residues L206-P212 and L207-S230 were replaced by a GGGG linker, respectively (Fig 5b). Astonishingly, the lid deletion variants were also functional in XopQ/Roq1 cell death assays (Fig 5c). The *Sl*EDS1 amino acid exchange variants and lid deletion variants accumulated to levels similar as wild type *Sl*EDS1 (Fig S5). These data suggest that if the *Sl*EDS1 catalytic triad residues mediate binding of a small molecule in *Nb* TNL immunity, perturbations of the triad environment and loss of the αF-helix lid are tolerated to a surprising extent to maintain *Sl*EDS1 immune functions.

**Figure 5:**
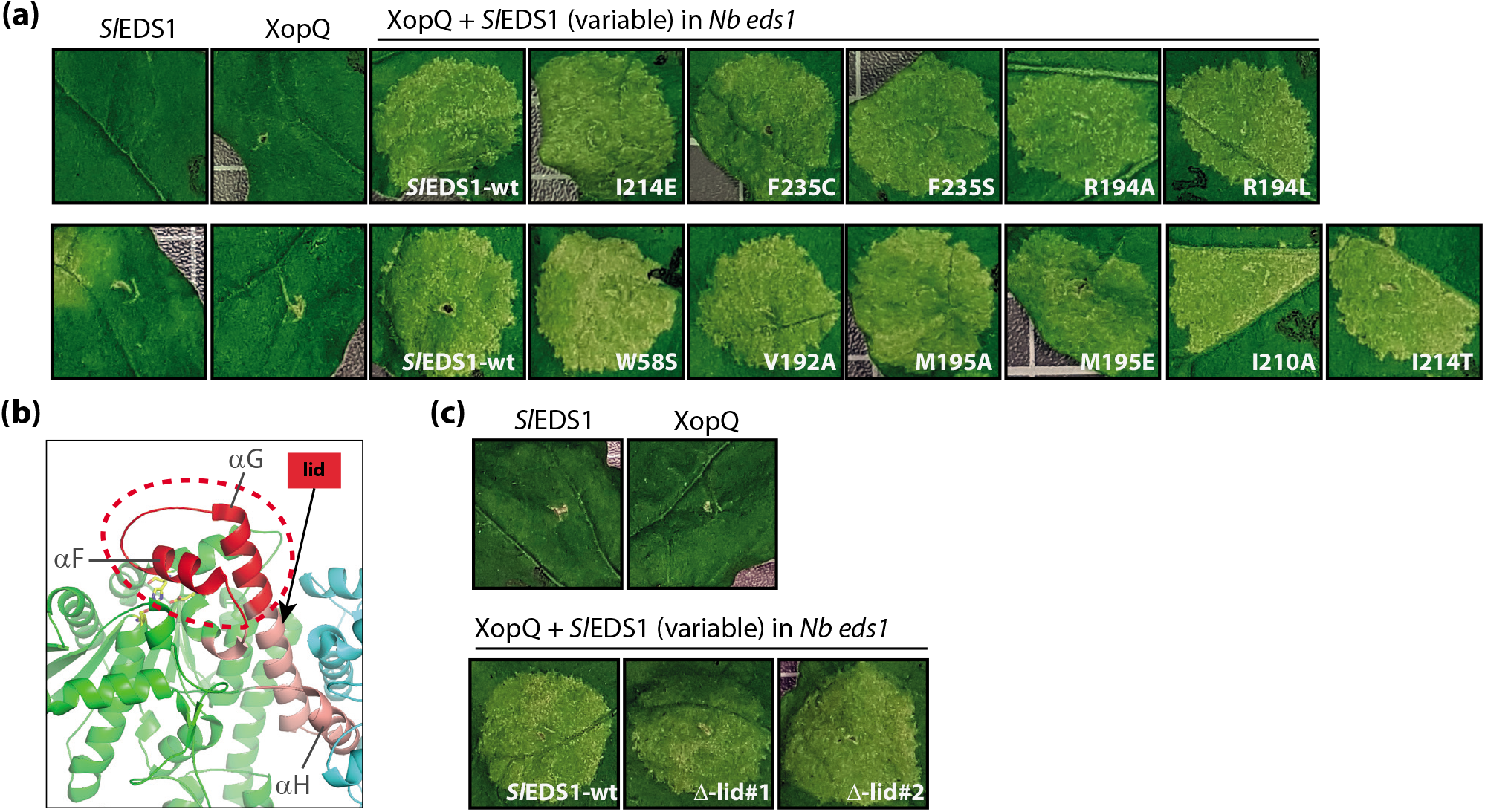
Perturbation of the *Sl*EDS1 catalytic triad environment. **(a)** Indicated single amino acid exchange variants of *Sl*EDS1 were co-expressed together with XopQ in *Nb eds1* mutant plants. XopQ/Roq1-mediated cell death was documented 6 dpi. The experiment was repeated at least five times with similar results, and representative images are shown. See also Fig S5. **(b)** Schematic representation of *Sl*EDS1 lid deletions. In two different constructs, the section marked in red, or parts thereof, was replaced by a GGGG linker sequence. green: α/ß hydrolase core fold, cyan: SAG101b, pink: lid region and αH helix, red: exposed lid region. **(c)** Lid deletions are functional in XopQ/Roq1 cell death assays. As in **(a)**, but lid deletions were tested in cell death assays.

### Nuclear EDS1 complexes are sufficient for XopQ/Roq1 cell death in *N. benthamiana*

We aimed to test in which subcellular compartment *Sl*EDS1-*Sl*SAG101b complex activity is required for *Nb* TNL signaling. We generated constructs for expression of *Sl*EDS1, *Sl*SAG101b or mis-localized variants, fused to Yellow Fluorescent Protein (YFP; Fig 6a). YFP-tagged *Sl*EDS1 or *Sl*SAG101b variants were transiently co-expressed with the respective mCherry-tagged complex partner and XopQ in *Nb epss* quadruple mutant plants deficient in all EDS1 family proteins. Subcellular localization and accumulation of proteins were monitored at 3 dpi by live cell imaging and immunodetection, respectively. Development of the XopQ/Roq1-induced cell death indicative of immune competence of *Sl*EDS1-*Sl*SAG101b was documented at 6 dpi.

**Figure 6:**
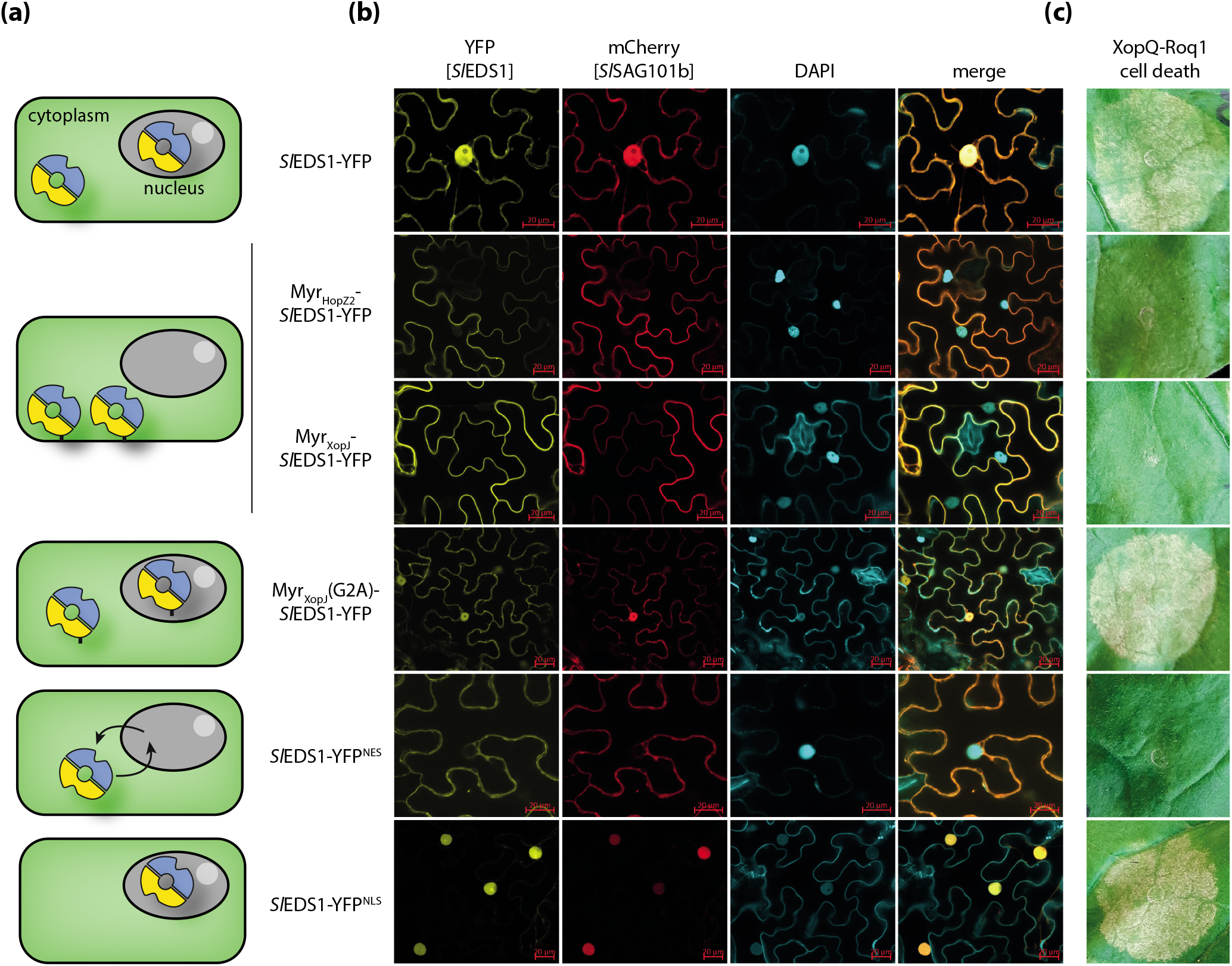
Immune-competence of mislocalized *Sl*EDS1 variants in *N. benthamiana* XopQ/Roq1 cell death assays. **(a)** Scheme of modifications for mislocalization of *Sl*EDS1 complexes. EDS1-YFP without modifications is detected in nucleus and cytoplasm (Gantner et al., 2019). Myristoylation motifs tether EDS1 and also SAG101b to the membrane. The G2A mutation abolishes myristoylation. NES leads to export from the nucleus, but does not prevent import, leading to import/export cycles. NLS facilitates enhanced import. **(b)** Live cell imaging of (mislocalized) *Sl*EDS1-*Sl*SAG101b expressed by âgroinfiltration in *Nb epss* plants. All images show single planes, and micrographs were taken 3 dpi. Localizations were determined for 3 independent experiments; representative micrographs are shown. Protein accumulation of variants is shown in Figure S6a. Scale bar = 20 μM. **(c)** Cell death signaling by mislocalized *Sl*EDS1-*Sl*SAG101b complexes. Induction of Roq1-mediated cell death upon co-expression of (mislocalized) *Sl*EDS1(YFP)-*Sl*SAG101b(mCherry) complexes with XopQ in *Nb epss* plants was documented 6 dpi. Cell death phenotypes were assessed in five independent experiments with similar results.

For mislocalization, the fusion proteins were either decorated with a nuclear localization signal (e.g., *Sl*EDS1-YFP^NLS^), a myristoylation motif (Myr-*Sl*EDS1-YFP) or a nuclear export signal (*Sl*EDS1-YFP^NES^). Attachment of a strong NLS was expected to deplete the cytosolic *Sl*EDS1-*Sl*SAG101b pool by enhancing nuclear import (Fig 6a; Garcia et al., 2010; Stuttmann et al., 2016). We used the classical SV40 NLS and that of c-myc, the human cancer protein. As similar results were obtained, only fusions containing the SV40-NLS are shown. The NES signal (from PKI; Wen et al., 1995) mediates enhanced nuclear export but does not prevent nuclear import, and thus promotes shuttling between nucleus and cytoplasm and increased cytosolic accumulation (Fig 6a; Garcia et al., 2010). Attachment of a myristoylation motif was conducted to tether complexes to the plasma membrane, thus preventing cytosolic movement and nuclear import (Fig 6a). Therefore, both NES and myristoylation motifs were expected to deplete nuclear protein pools. We used the myristoylation motifs of the effectors HopZ2 and XopJ (Myr_HopZ2_ and Myr_XopJ_), and generated G2A variants lacking a critical glycine residue as control (Lewis et al., 2008; Bartetzko et al., 2009). We present only data using Myr_XopJ_(G2A) constructs as similar results were obtained with both motifs. In two sets of experiments, *Sl*EDS1 (Fig 6) or *Sl*SAG101b (Fig S6) YFP fusions were mislocalized. Because stronger plant responses were obtained with the *Sl*EDS1-YFP fusions, we continued assays with *Sl*EDS1-YFP mis-localized forms.

As expected, transiently co-expressed *Sl*EDS1-YFP and *Sl*SAG101b-mCherry complexes localized to the nucleus and cytoplasm (Fig 6b; Gantner et al., 2019), and were functional in XopQ/Roq1 cell death assays (Fig 6c). When *Sl*EDS1-YFP^NES^ or fusions containing functional myristoylation motifs were expressed with *Sl*SAG101b-YFP, nuclear pools were diminished, as both *Sl*EDS1 and *Sl*SAG101b were detected mainly in the cytoplasm and nuclear periphery (NES) or at the plasma membrane (Myr; Fig 6b). A G2A mutation in Myr_XopJ_(G2A)-*Sl*EDS1-YFP restored nucleo-cytoplasmic distribution of *Sl*EDS1 and *Sl*SAG101b (Fig 6b). Indeed, strongly reduced XopQ/Roq1 cell death was detected upon depletion of the nuclear *Sl*EDS1-*Sl*SAG101b pool, and was restored upon co-expression of the Myr_XopJ_(G2A)-S*l*EDS1-YFP variant. In contrast, expression of EDS1-YFP^NLS^ resulted in detection of *Sl*EDS1-*Sl*SAG101b predominantly in the nucleus (Fig 6b) and wild type-like XopQ/Roq1 cell death induction (Fig 6c). These results agree with earlier indications in Arabidopsis (Garcia et al., 2010; Stuttmann et al., 2016) that nuclear EDS1 complexes are required and sufficient for TNL immunity. Nevertheless, we cannot exclude the possibility that low accumulation of the EDS1-YFP^NES^ fusion (Fig S6) or membrane tethering of Myr-*Sl*EDS1-YFP might interfere with protein function, beyond simply reducing its accumulation in the nuclear compartment.

## Discussion

EDS1 complexes, together with ADR1- and NRG1-type hNLRs, form immunity signaling modules which in Arabidopsis are essential for TNL-initiated ETI and contribute to CNL-ETI, PTI and basal immunity (Dongus and Parker, 2021). We demonstrate in *Nb* that EDS1-RNL modules are dispensable for several tested PTI responses (Fig 1). Therefore, the role of EDS1 in complex with SAG101b appears to be more aligned with signal relay from activated receptors to RNLs in TNL immunity. Intriguingly, exchanges of lipase-like domain catalytic triad residues in immune-competent *Sl*EDS1 can enhance or abolish TNL immunity in *Nb* (Fig 2), in support of a small molecule binding to this region and contributing to the immune response. Further *Sl*EDS1 mutagenesis and heterologous assays suggest that if the EDS1 or PAD4 α/β-hydrolase pocket has ligand-binding activity, it is variable between Arabidopsis and *S. lycopersicum* orthologs and not easily perturbed by single amino acid changes in the *Sl*EDS1 catalytic pocket environment or lid (Figs 3-5). The occurrence of a degenerate lipase pocket in SAG101 orthologs is consistent with EDS1-SAG101 dimers operating *via* their conserved EP-domains rather than using the lipase triad in TNL immunity (Gantner et al., 2019; Lapin et al., 2019). By contrast, the observed difference in importance of triad residues between *At* and *Sl* EDS1 proteins (Figs 2,3), as well as a requirement for an intact *At*PAD4 lipase pocket in resistance to green peach aphid feeding (Louis et al., 2012; Dongus et al., 2020), leaves open the possibility that certain EDS1 and PAD4 protein states utilize this domain for ligand interaction, as observed for a number of plant lipase-like hormone receptors (Mindrebo et al., 2016). Our data further support an important function of *Sl*EDS1-*Sl*SAG101b complexes within the nucleus (Fig 6). This prompts the question how and in which compartment RNLs are activated *via* EDS1 dimers to form proposed membrane pores.

### *N. benthamiana* LRR-RLK and LRR-RLP-mediated PTI and ETI-PTI connectivity

A dual role in plant development and immunity for LRR-RLKs from the SERK family was firmly established in different species (Chinchilla et al., 2007; Fradin et al., 2009; Chen et al., 2014). In *Nb*, virus-induced gene silencing of *BAK1/SERK3*-like genes led to a strong reduction of the flg22-induced ROS burst response (measured with 50 nM flg22; Heese et al., 2007). The *Nb bak1* mutant generated in this study displayed a reduced ROS burst when challenged with 2 nM flg22 (Figs 1, S1-3), but was indistinguishable from wild type with a higher ligand concentration (50 nM flg22). Similarly, reduced Cf4/Cf9-mediated cell death (Figs 1, S3) only became apparent when low inoculum densities were used for agroinfiltration. These differences suggest that additional *SERK* family genes, silenced by VIGS but not inactivated in the *Nb bak1* double mutant, contribute to *Nb* PTI signaling. Indeed, a recent chromosome-level assembly of the *Nb* genome (https://www.nbenth.com/; v3.5) encodes four protein orthologs most similar to *At*BAK1 in reciprocal BLAST searches (Table S1). The respective mRNAs have extensive similarity to the silencing construct used by Heese et al. (2007). It is therefore plausible that the two further *BAK1-like* genes that were not targeted by CRISPR/Cas partially mediate LRR-RLK and LRR-RLP responses in the *Nb bak1* mutant line. This is also supported by the mild stunting of *NbBAK1*-VIGS plants (Heese et al., 2007) but not the *Nb bak1* line used in this study. Further *SERK* family genes might also contribute to *Nb* PTI signaling, as previously suggested (Fradin et al., 2011; Postma et al., 2016).

PTI assays including mutant *Nb* lines deficient in EDS1 family proteins or RNLs were conducted with low inoculum densities (Avr4/9) and elicitor concentrations (flg22). These assay conditions allowed detection of weakly impaired PTI in *Nb bak1*, whereas *eds1* or *rnl* mutant lines behaved like wild type in the same experiments (Figs 1, S3). These results suggest that RLP- and RLK-mediated PTI responses, as far as tested, do not require an intact TNL signaling sector in *Nb*. Recruitment of EDS1-RNL modules for PTI signaling (Pruitt et al., 2021; Tian et al., 2021) might thus not represent a conserved function of these TNL immune sector components. Alternatively, EDS1-RNL modules in *Nb* could support signaling by surface receptors not analyzed here and/or promote a different set of signaling outputs.

Helper NLRs of the NRC (NLR Required for Cell Death) superclade were found to be required for cell death initiated by Cf4/Avr4 and further LRR-RPs in *Nb* (Gabriels et al., 2006; Gabriels et al., 2007; Fradin et al., 2009). Analysis of various CRISPR/Cas-induced *Nb nrc* mutant lines determined that in fact NRC3 confers Cf4-initiated cell death (Kourelis et al., 2021). Therefore, dependency of PTI responses on intracellular immune receptor networks might be a conserved property but executed by diverse mechanisms for different receptors or particular plant species, in agreement with large structural and functional diversity observed among both surface-localized as well as intracellular immune receptors (Zipfel, 2014; Van de Weyer et al., 2019; Lu and Tsuda, 2021; Pruitt et al., 2021). Notably, PTI-ETI connections might correlate with evolutionary trajectories and expansions of specific immune sectors in species: Whereas Arabidopsis has a large number of TNLs, *Nb* has relatively few TNLs but expanded NRCs (Hofberger et al., 2014; Wu et al., 2017; Johanndrees et al., 2021).

### Role of EDS1 complexes in TNL immunity signaling

A core function of EDS1 complexes is in signal transmission from activated TNL-type immune receptors to RNL-type hNLRs. NADase activity has been reported for a number of plant TIR domains, full length TNLs RPP1 and Roq1 and TIR domain-containing proteins from bacterial and animal origins (Essuman et al., 2017; Essuman et al., 2018; Horsefield et al., 2019; Wan et al., 2019; Ma et al., 2020; Martin et al., 2020; Ofir et al., 2021). Additionally, plant individual TIR domains appear to also function as 2′,3′-cAMP/cGMP synthetases, an activity required together with the NADase activity for mediating TNL immunity-related cell death (Yu et al., 2021). In animal axon degeneration, depletion of NAD^+^ by TIR-domain protein SARM1 (sterile alpha and Toll/interleukin-1 receptor motif-containing 1) initiates a neuronal self-destruction program (Essuman et al., 2017). A comparable mechanism is unlikely to account for plant TNL signaling, since plant TIR domains demonstrate comparably low NADase activity and do not deplete cellular NAD^+^ stores. Also, NAD^+^ depletion is not sufficient for induction of TNL immunity (Wan et al., 2019; Duxbury et al., 2020). Thus, a mechanism in which one or several products of TIR enzymatic activities function as signal intermediates is more likely in TNL immunity.

There are two candidate regions for small molecule binding to EDS1 complexes. One is a cavity produced by the EDS1 heterodimers and lined by several conserved and essential positively charged residues (Wagner et al., 2013; Bhandari et al., 2019). The second is the lipase-like domain with a conserved catalytic triad and assumed substrate binding site in EDS1 and PAD4 proteins. A role for the PAD4 catalytic triad was previously uncovered in Arabidopsis resistance to green peach aphid (GPA; Louis et al., 2012). The PAD4 lipase-like domain, in absence of EDS1, is sufficient for full GPA resistance and depends on the triad serine and aspartate, but not histidine, residues (Louis et al., 2012; Dongus et al., 2020).

Abolished or enhanced immune functions of *Sl*EDS1 triad mutants in *Nb* (Fig 2) provide a first hint that the EDS1 enzyme signature might also be of relevance for TNL immunity. Functionality of the *Sl*EDS1^S125A^ variant suggests that catalysis of a potential substrate is not required for TNL immunity. This could point towards a mechanism as proposed for the α/ß-hydrolase receptor D14 (Dwarf14) in strigolactone perception (Seto et al., 2019): In this receptor system, the hormone first binds the D14 receptor, which requires the catalytic triad serine and histidine residues. Hormone binding induces a conformational change and downstream signaling, which depends on structural reorganization of the lid domain and interactor recruitment. Subsequently, hydrolytic cleavage of the hormone, following slow kinetics, deactivates signaling. Rice GID1 is another example of a plant α/ß-hydrolase functioning as a small molecule receptor. GID1 lacks the triad histidine residue and has no catalytic activity. In GA signaling, the GID1 N-terminal lid domain wraps over bound GA and forms a new surface for recruitment of DELLA proteins (Mindrebo et al., 2016).

Although a mechanistic analogy is tempting, several pieces of evidence argue against binding of a small molecule to the *Sl*EDS1 lipase-like domain active site region for activation of downstream RNLs. Deduced from the *At*EDS1-*At*SAG101 crystal structure, *At*EDS1 has features of both active and inactive α/ß-hydrolases (Wagner et al., 2013; Voss et al., 2019). A perturbed oxyanion hole would be in agreement with potentially slow substrate catalysis for inactivation of signaling, as described for the D14 strigolactone receptor (Seto et al., 2019). Also, the αF lid helix, which shields the presumed substrate binding pocket in the crystal structure, might assume an alternative conformation upon ligand binding. However, an appreciable cavity for binding of a larger substrate was not detected (Wagner et al., 2013; Voss et al., 2019). Here, *Sl*EDS1 immune functions were not disturbed by any of the additional exchanges that were introduced in proximity of the catalytic triad residues (Fig 5). It is possible that key residues were missed or combined exchanges required to disrupt binding, but this would imply a remarkably robust receptor-ligand interaction. Also deletion of the entire lid region did not abolish XopQ/Roq1 cell death in complementation assays (Fig 5). Furthermore, we did not detect significant impairment of immune functions in catalytic triad variants of *Sl*EDS1-*Sl*PAD4 and *At*EDS1-*At*PAD4 in Arabidopsis, or *At*EDS1-*At*SAG101 in *Nb* (Figs 3,4). Considering all of these features, our data do not conclusively support a function of the EDS1 active site region involving small molecule binding in TNL immunity.

### Subcellular compartmentalization of TNL immune signal relay in *N. benthamiana*

Current TNL activation and signaling models invoke small molecule production *via* TIR-NADase and/or TIR-2′,3′-cAMP/cGMP synthetase activities (Horsefield et al., 2019; Wan et al., 2019; Yu et al., 2021). TIR-derived small molecules can be expected to be mobile within the cell, and could therefore be produced in different subcellular compartments in alignment with immune receptor localization and function. Indeed, subcellular localizations including nucleus, plasma membrane and endomembrane systems were reported for TNLs (e.g., Takemoto et al., 2012). However, at least the TNLs N (from *N. tabacum*) and RPS4 (from Arabidopsis and functioning in a pair together with RRS1; (Narusaka et al., 2009; Saucet et al., 2015)) localize within the nucleus for effective TNL immunity (Burch-Smith et al., 2007; Wirthmueller et al., 2007; Huh et al., 2017). Similarly, nuclear EDS1 complexes are sufficient for natural pathogen resistance and TNL autoimmunity in Arabidopsis, while nuclear exclusion impairs TNL immune responses (Garcia et al., 2010; Stuttmann et al., 2016; Ordon et al., 2021). The subcellular localization of *Nb* Roq1 and site of XopQ recognition was so far not reported. Results presented here from mis-localizing *Sl*EDS1-*Sl*SAG101b complexes suggest that activation of EDS1 complexes inside nuclei is sufficient and also critical for Roq1 cell death signaling in *Nb* (Fig 6).

Activated forms of RNL-type hNLRs NRG1 and ADR1, functioning downstream or together with EDS1 complexes in Arabidopsis and *Nb*, were reported to localize to the plasma membrane, endoplasmatic reticulum membrane and the cytosol (Wu et al., 2018; Lapin et al., 2019; Jacob et al., 2021; Saile et al., 2021). In the case of *At*ADR1, counteracting plasma membrane localization by depletion of phosphatidylinositol-4-phosphate impaired ADR1-mediated cell death induction, further supporting membrane-associated functions of ADR1 in immunity (Saile et al., 2021). Nuclear localization or functions of RNLs were so far not reported. It is thus of major interest for future analyses to determine how and in which compartment EDS1-RNL interactions (Qi et al., 2018; Sun et al., 2021; Wu et al., 2021) mediate RNL activation and TNL immunity.

## Materials and methods

### Plant material and growth conditions

*Nb* lines used were *Nb eds1a-1* (Ordon et al., 2017), *pad4-1 sag101a-1 sag101b-1* (Gantner et al., 2019), *nrg1-4* (Ordon et al., 2021) and *eds1 pad4 sag101a sag101b* (*epss*; Lapin et al., 2019). The *Nb bak1* mutant line was generated by CRISPR/Cas using previously described constructs (Stuttmann et al., 2021), and additional details are provided in Fig S1 and Table S1. The *Nb adr1 nrg1* double mutant line is described in more detail elsewhere (Prautsch et al., 2021). *Nb* plants were cultivated in a greenhouse with a 16-h light period (sunlight and/or IP65 lamps [Philips] equipped with Agro 400 W bulbs [SON-T]; 130–150 μE/m^−2^ s^−1^; switchpoint; 100 μE/m^−2^ s^−1^), 60% relative humidity at 24/20°C (day/night). Arabidopsis accession Columbia-0 was used, and plants were cultivated under short-day conditions (8 h light, 23/21°C [day/night], 60% relative humidity) or in a greenhouse under long-day conditions (16 h light) for seed set. An *eds1-12 pad4-1 sag101-3* triple mutant line was generated by crossing the *eds1-12* line (Ordon et al., 2017) with a *pad4-1 sag101-3* double mutant line (Cui et al., 2018).

### Molecular cloning and plant transformation

Gateway cloning and Golden Gate assembly were used to generate plant expression constructs using the Modular Cloning Plant Toolbox, Plant Parts I and II parts collections (Engler et al., 2014; Gantner et al., 2018). Newly generated plasmids, including Level 0 modules, and oligonucleotides used in this study, are described in Tables S2 and S3, respectively. ccdB survival II cells (Thermo Fisher) and Dh10b/Top10 cells were used for vector propagation and cloning, *Agrobacterium tumefaciens* strain GV3101 pMP90 was used for transient expression and stable plant transformation. Arabidopsis plants were transformed by floral dipping as previously described (Logemann et al., 2006). *Nb* transformation was conducted following our protocol provided on the protocols.io platform (https://doi.org/10.17504/protocols.io.sbaeaie).

### Transient expression, infection and ion leakage assays

For transient protein expression in *Nb* (agroinfiltration), plate-grown Agrobacteria were resuspended in Agrobacterium Infiltration Medium (10 mM MES pH 5.8, 10 mM MgCl_2_) and infiltrated with a needleless syringe at an OD_600_ = 0.4 per strain, if not indicated otherwise. To measure ion leakage, leaf discs were harvested using a biopsy punch (5 mm) into 24 well plates. Leaf discs were washed 2h in H_2_O under mild agitation, H_2_O was replaced and conductivity was measured 24 hours later using a handheld conductivity meter (LAQUAtwin COND, Horiba Scientific). *Xanthomonas campestris* pv. *vesicatoria* strain 85-10 (Thieme et al., 2005) was used for mixed co-infiltrations and resistance assays, as previously described (Lapin et al., 2019). *Pseudomonas syringae* pv *tomato* infection assays were conducted as previously described (Lapin et al., 2019). For infections with *Hyaloperonospora arabidopsidis* (*Hpa*), T_1_ transgenic seeds were selected by fluorescence (FAST; Shimada et al., 2010). Plants were grown under short day conditions for 3 weeks, infected with *Hpa* isolate Cala2 and tissues used for Trypan Blue staining at 5 – 7 dpi, as previously described (Stuttmann et al., 2011). Red light imaging of cell death (Landeo Villanueva et al., 2021) was conducted using a Vilber Fusion FX system. Intensity of cell death reactions was measured using ImageJ, and data were normalized to a 0-1 range.

### Immunodetection, protein co-purification and live cell imaging

Proteins were extracted by direct grinding of tissues in Laemmli buffer, denaturation, and clearing by centrifugation. Extracts were resolved on 8 – 12% SDS-PAGE gels, and transferred to nitrocellulose membranes (GE Healthcare) for immunodetection. Membranes were stained with Ponceau or Amido black to control loading as described (Goldman et al., 2016). Strep-Tactin-AP conjugate (IBA GmbH) was used for detection of Strep-tagged proteins. Primary antibodies used were α-GFP (mouse) and α-hemagglutinin (rat; both from Roche). Horseradish peroxidase-coupled (GE Healthcare) or alkaline phosphatase-coupled (Sigma-Aldrich, www.sigmaaldrich.com) secondary antibodies were used. A Zeiss LSM780 confocal laser scanning microscope was used for live cell imaging. All images are single planes. DAPI staining was used to mark nuclei.

### PTI assays: ROS measurements

Production of reactive oxygen species (H_2_O_2_) was measured in a 96-well plate format using leaf discs of 4-week-old *Nb* plants as previously described (Gomez-Gomez et al., 1999). Briefly, leaf discs were cut with a 5 mm biopsy punch (PFM Medical), elicited with 2 nM flg22 (synthesized in-house) and measured in 2 min intervals (60 min total time) with a TriStar2 S LB942 luminescence plate reader (Berthold Technologies).

## Acknowledgements

This work was funded by GRC grant STU 642-1/1 (Deutsche Forschungsgemeinschaft, DFG) to JS. Furthermore, JS is grateful for support through the Leibniz price from the DFG and the Alfried Krupp von Bohlen und Halbach Stiftung, awarded to Ulla Bonas. DL and JEP acknowledge support from The Max-Planck Society and Deutsche Forschungsgemeinschaft (DFG, German Research Foundation) SFB-1403–414786233. We are grateful to Bianca Rosinsky for taking care of plant growth facilities and growing plants. We thank Bart Thomma for providing Avr4/Cf4 and Avr9/Cf9 expression constructs and Sebastian Schornack for discussion on BAK1 orthologs.

## Author contributions

JZ, JG, DL, KB, LEL, SZ, and CK performed experiments and analyzed data. JZ, DL, KB and LEL contributed to figure preparation. RG performed molecular modeling and structural analyses. JS conceptualized and supervised work, performed experiments and analyzed data, prepared figures, and wrote the manuscript with contributions from DL and JEP. All authors approved the final version.

---

**Figure S1:**
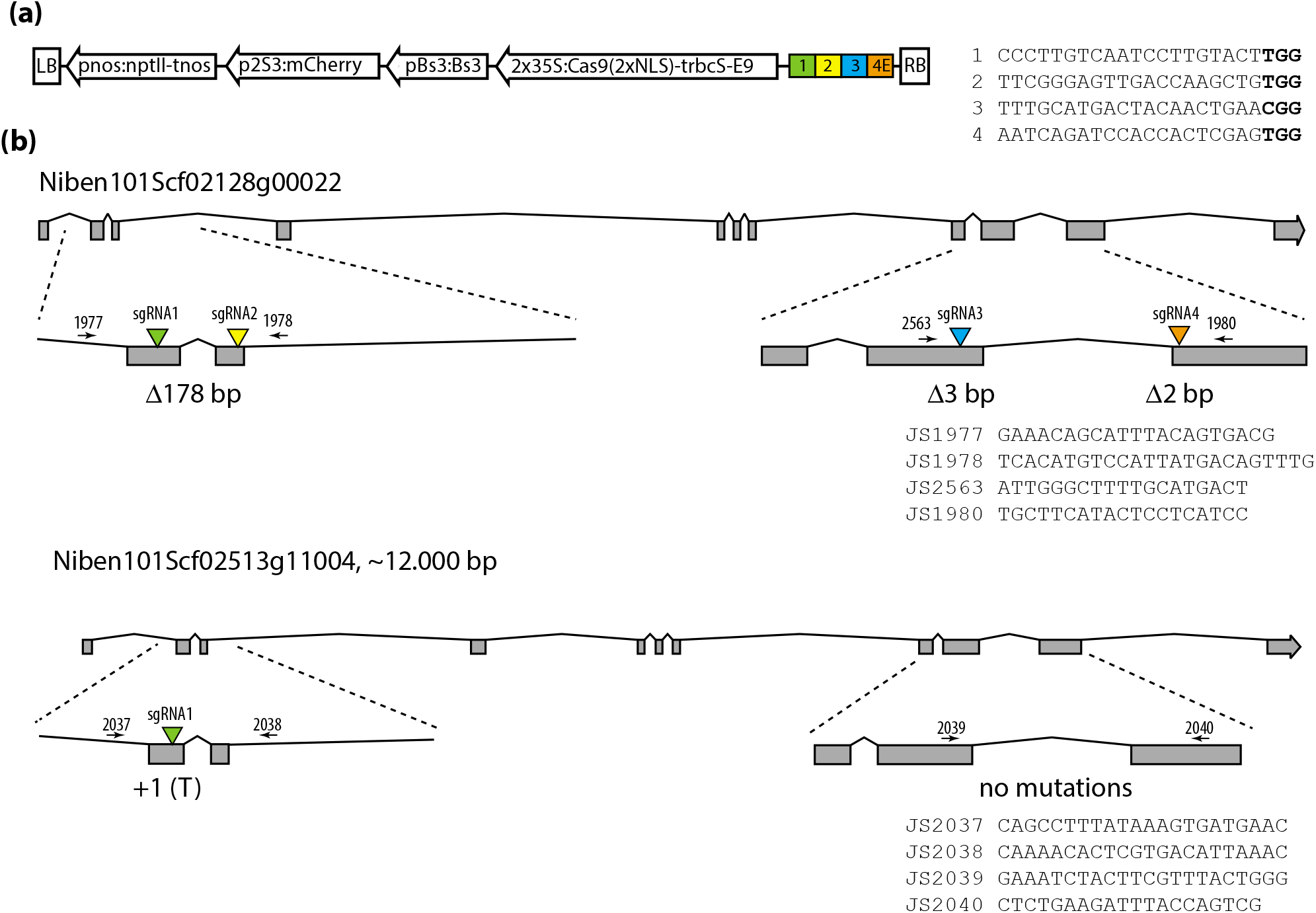
Generation of an *N. benthamiana bak1/serk3* mutant line by genome editing. **(a)** Scheme of pDGE492 used for generation of *Nb bak1* mutant plants; based on pDGE463 (Stuttmann et al., 2021). Guide RNAs were expressed under control of a tomato U3 promoter fragment (Stuttmann et al., 2021). Target sites are depicted, color code corresponds to panel b. **(b)** Gene models of *BAK1/SERK3*-like genes targeted for editing. Mutations detected in the line used in this study are depicted below the gene model, and primers used for genotyping are shown. The *Nb* genome assembly (www.nbenth.com) encodes four proteins most similar to *At*BAK1 from Arabidopsis. Only the two genes shown here were edited in *Nb bak1*, as verified by Sanger sequencing of PCR products derived from all four genes. Additional details on gene models are provided in Table S1.

**Figure S2:**
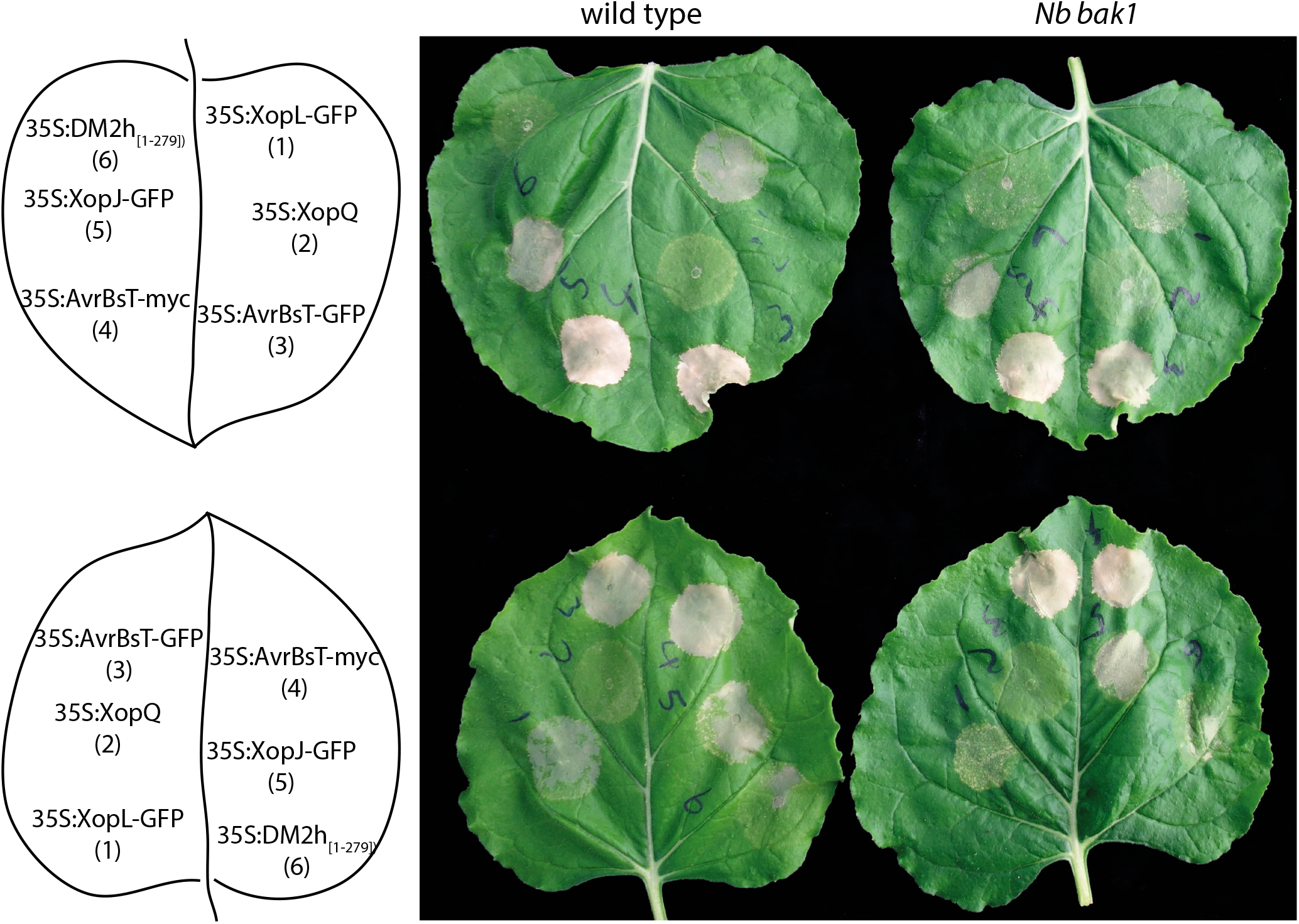
*N. benthamiana bak1* mutant plants are not impaired in cell death responses induced by intracellular pathways. Constructs with depicted T-DNA cassettes were used for transient expression (by agroinfiltration). Development of the hypersenitive response was monitored daily, and documented 4 dpi. Similar HR development between wild-type and *Nb bak1* mutant plants was observed in four independent experiments, each including at least four leaves and two independent plants of each genotype. XopJ, XopL and AvrBsT are effectors from *Xanthomonas campestris* pv. *vesicatoria* (*Xcv*) that induce cell death by yet unknown mechanisms. *Xcv* XopQ is recognized by the TNL Roq1 (Schultink et al., 2017). DM2h_(1-279)_ is an autoactive fragment of the DM2h TNL receptor from Arabidopsis (Ordon et al., 2021).

**Figure S3:**
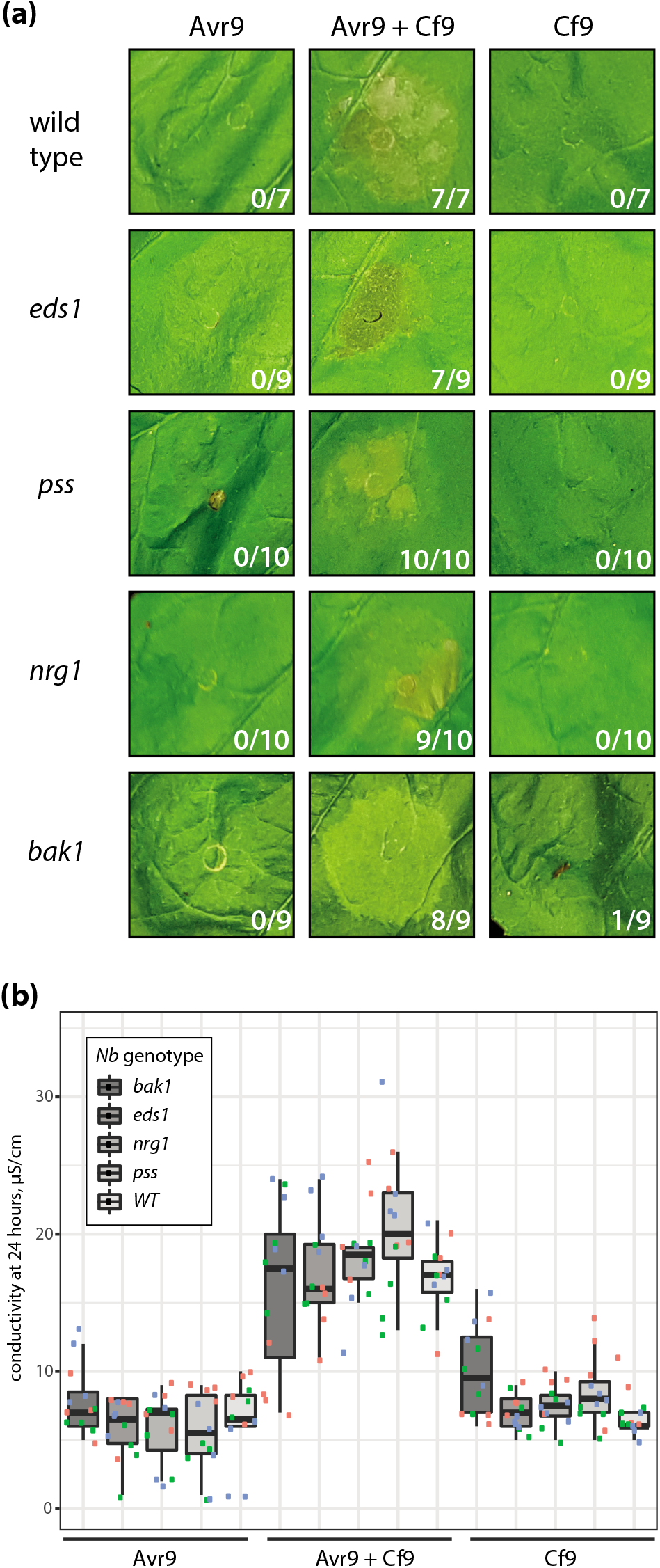
Cf-9/Avr9-induced cell death in *EDS1* family and *RNL* mutant *N. benthamiana* lines. **(a)** Cell death induction upon (co-)expression of Avr9 and Cf-9. Avr9 and/or Cf-9 were expressed by agroinfiltration (OD_600_ = 0.1 (Cf9), 0.2 (Avr9)) in the indicated *Nb* lines. Symptom (cell death) formation was documented 4 dpi. The experiment was conducted four times with similar results, and representative images are shown. Numbers indicate infiltration sites with chlorosis (*bak1*) or cell death (all remaining genotypes). **(b)** As in (a), but ion leakage was measured 24 hpi. Individual data points from three independent experiments with four replicates are shown. Differences within treatment groups were not statistically different (ANOVA).

**Figure S4:**
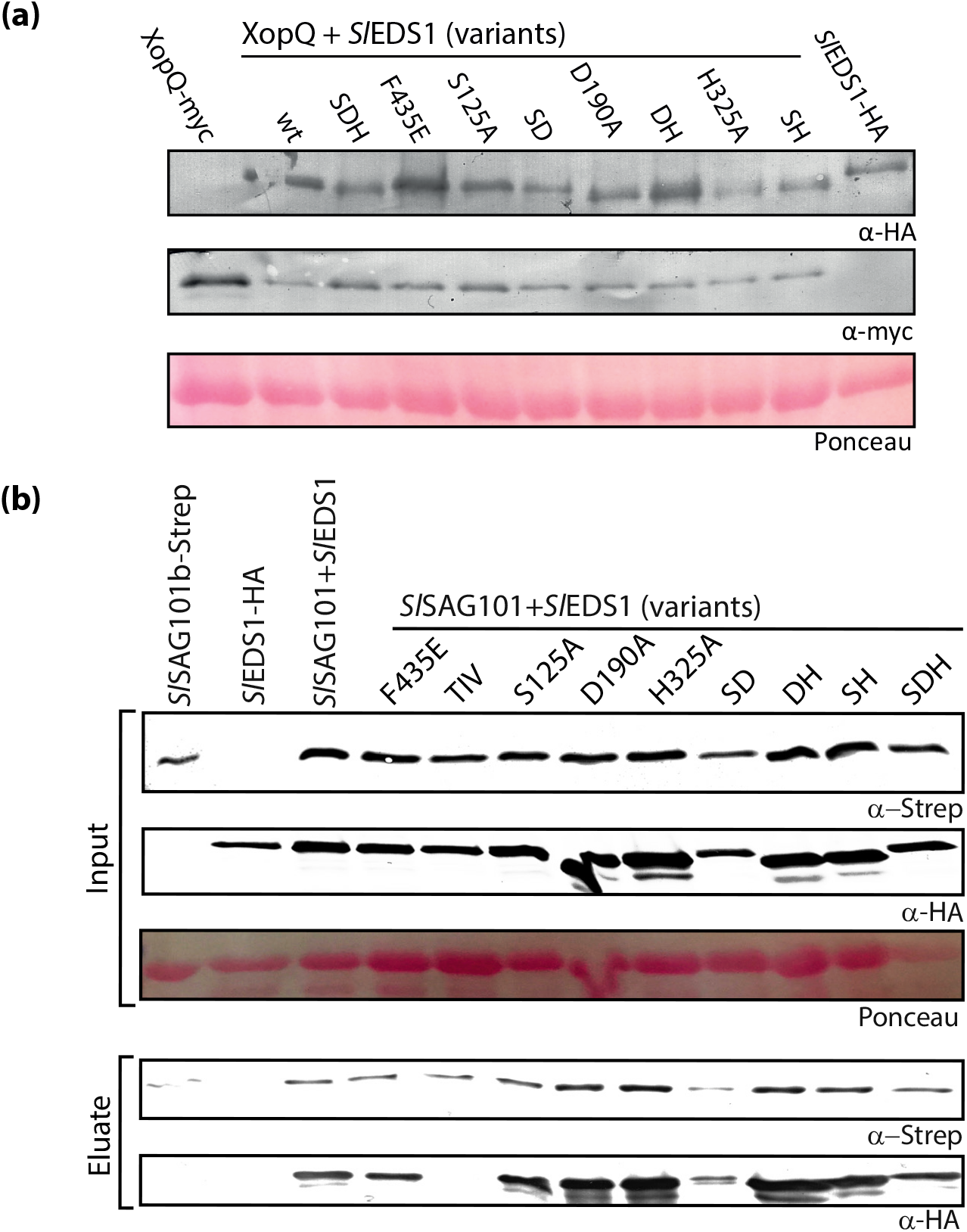
Stability of *Sl*EDS1 catalytic triad variants, and complex formation with *Sl*SAG101b. **(a)** Immunodetection of proteins transiently expressed for cell death assays (Fig. 2). Samples were taken 3 dpi, protein extracts prepared by grinding tissues in Laemmli buffer and analysed by SDS-PAGE and immunodetection. Poinceau staining is shown as loading control. Immunodetection was included in several replicates of cell death assays, with similar results. **(b)** Co-purification assay with *Sl*SAG101b-Strep and *Sl*EDS1-HA or variants thereof. Indicated proteins were, alone or in combination, transiently expressed in *Nb*. Tissues were used for co-purifiaction 3 dpi, and input and eluate fractions were analyzed by SDS-PAGE and immunodetection. Co-purification assays including *Sl*EDS1 S/D/H single mutant and double mutant variants were conducted three times, with similar results.

**Figure S5:**
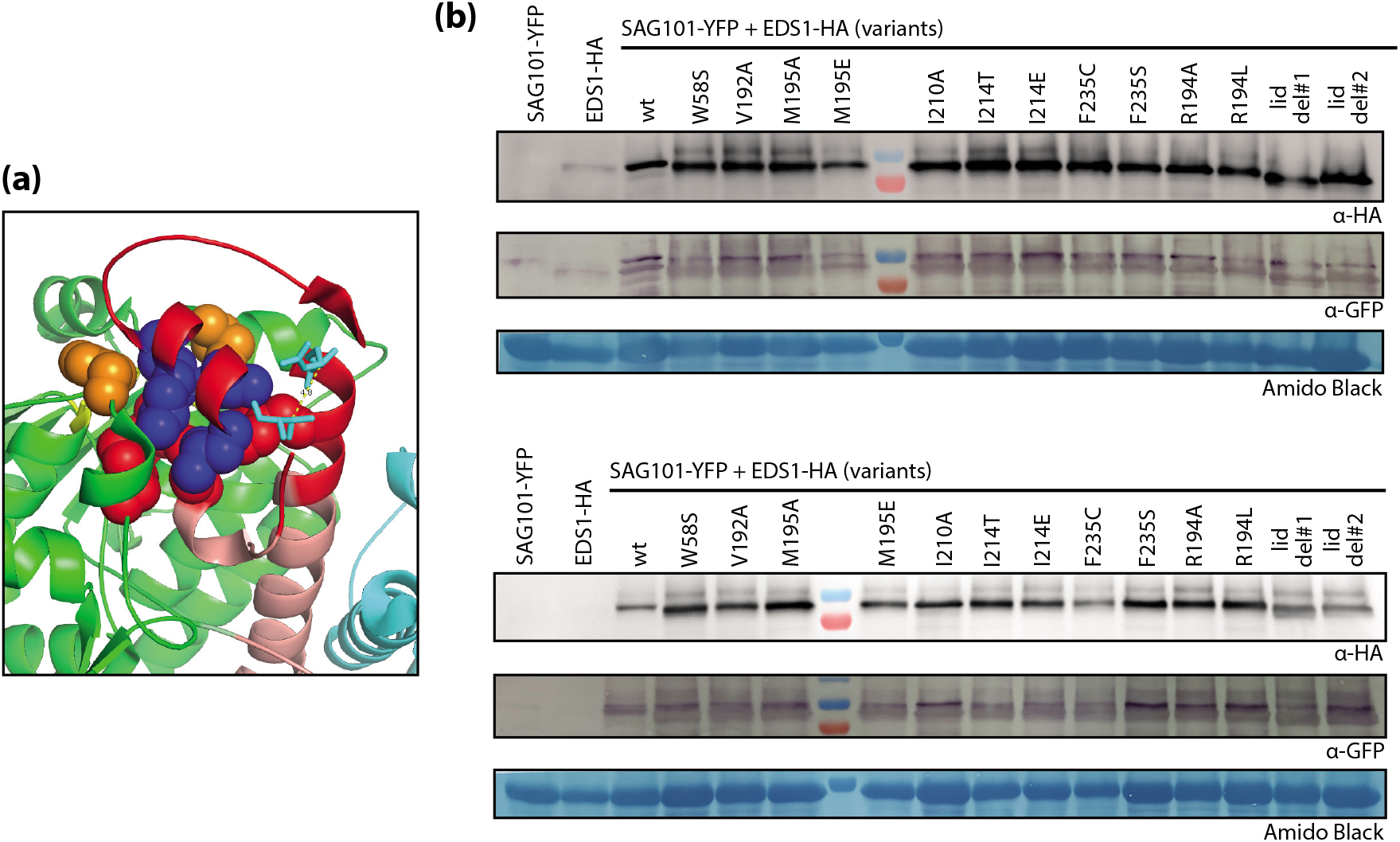
Immunodetection of *Sl*EDS1 lid deletion variants and exchanges in proximity of the catalytic triad. **(a)** Structural view of region targeted for perturbation of the triad environment. The *Sl*EDS1 lid region (red) with the αH helix (pink) and the core α/ß-hydrolase domain (green) is illustrated. Amino acids targeted for mutagenesis (see Figure 5a for functional data) are highlighted by spheres. **(b)** Protein accumulation of *Sl*EDS1 variants. Cell death assays (co-infiltrations with XopQ-myc, Figure 5a) were conducted in parallel, but *Sl*EDS1 variants (with HA tag) were co-expressed with *Sl*SAG101b-GFP (without XopQ) for immunodetection. Wild-type *Nb* were used for agroinfiltration, and tissues were harvested 3 dpi for preparation of samples for SDS-PAGE and western blotting. The lower section of the membrane was stained with Amido Black as a loading control. The same membrane was used to detect HA-tagged *Sl*EDS1 variants (by chemoluminescence), and subsequently *Sl*SAG101b-GFP (by alkaline phosphatase detection).

**Figure S6:**
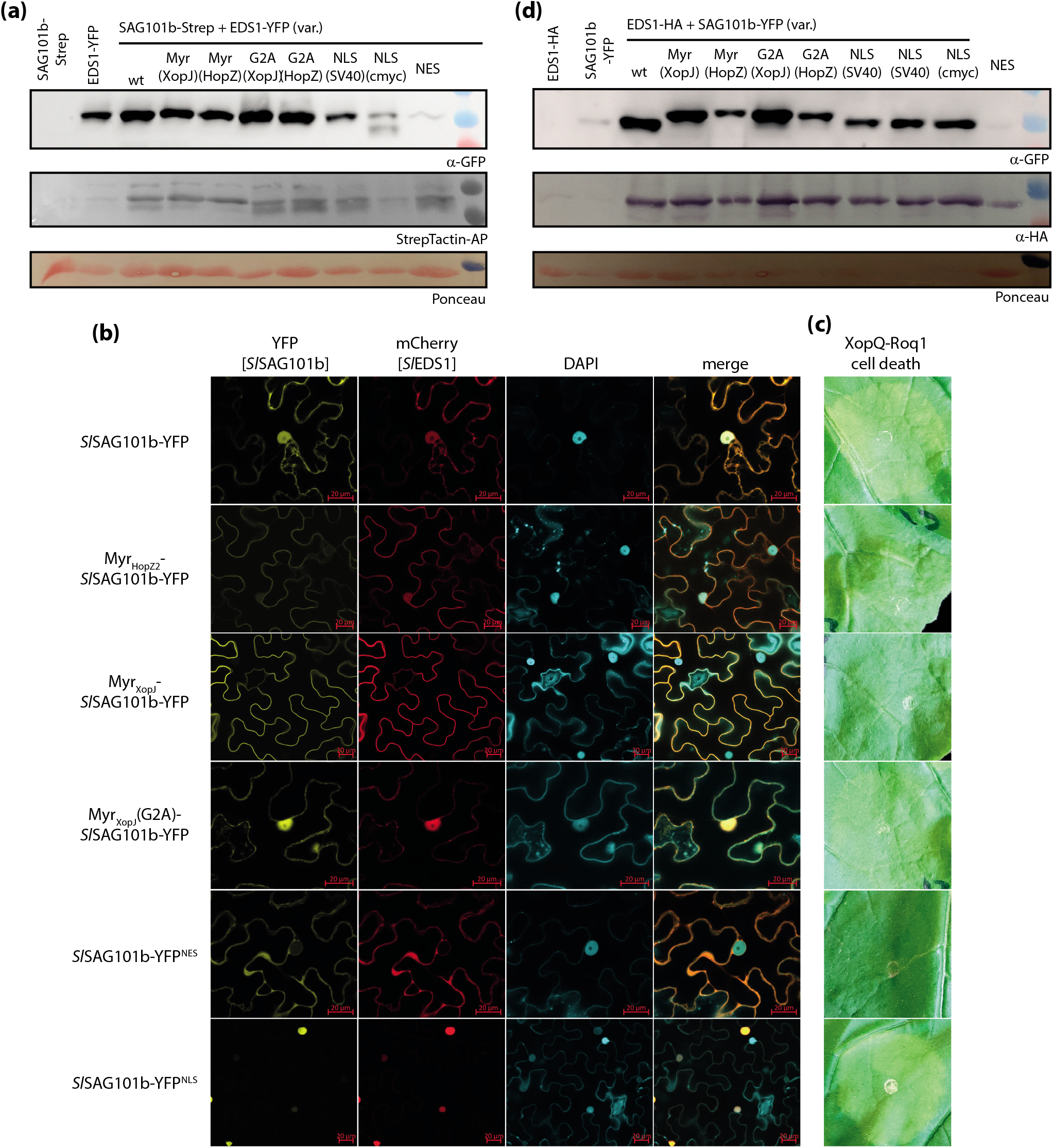
Immune-competence of mislocalized *Sl*SAG101b and protein accumulation of *Sl*EDS1 and *Sl*SAG101b mislocalization variants. **(a)** Accumulation of *Sl*EDS1-YFP and mislocalization variants in *Nb epss* tissues. *Sl*EDS1-YFP fusions were co-expressed together with *Sl*SAG101b-Strep to avoid interference from *Sl*SAG101b-mCherry (used for live cell imaging and cell death assays) during immunodetection. Protein samples were taken 3 dpi. Immunodetection was conducted 4 times. The NES variant consistently accumulated to low levels. **(b)** Live cell imaging of (mislocalized) *Sl*SAG101b-YFP and *Sl*EDS1-mCherry proteins expressed by agroinfiltration in *Nb epss* plants. All images show single planes, and micrographs were taken 3 dpi. Protein accumulation of variants is shown in Figure S6d. Scale bar = 20 μM. **(c)** Cell death signaling by mislocalized *Sl*EDS1(mCherry)-*Sl*SAG101b(YFP) complexes. As in (b), but cell death reactions were documented 6 dpi. Cell death assays were conducted 4 times with similar results; representative images are shown. **(a)** Protein accumulation of *Sl*SAG101b variants used in (b) and (c); in co-expression with *Sl*EDS1-HA (see also panel (a)). Immunodetection was conducted 3 times. The NES variant consistently accumulated to low levels.

## References

Adlung, N., Prochaska, H., Thieme, S., Banik, A., Bluher, D., John, P., Nagel, O., Schulze, S., Gantner, J., Delker, C., Stuttmann, J., and Bonas, U. (2016). Non-host Resistance Induced by the Xanthomonas Effector XopQ Is Widespread within the Genus *Nicotiana* and Functionally Depends on *EDS1*. Front Plant Sci 7, 1796.

Baggs, E.L., Monroe, J.G., Thanki, A.S., O’Grady, R., Schudoma, C., Haerty, W., and Krasileva, K.V. (2020). Convergent Loss of an EDS1/PAD4 Signaling Pathway in Several Plant Lineages Reveals Coevolved Components of Plant Immunity and Drought Response. The Plant Cell 32, 2158–2177.

Barragan, A.C., and Weigel, D. (2021). Plant NLR diversity: the known unknowns of pan-NLRomes. The Plant Cell 33, 814–831.

Bartetzko, V., Sonnewald, S., Vogel, F., Hartner, K., Stadler, R., Hammes, U.Z., and Bornke, F. (2009). The *Xanthomonas campestris* pv. *vesicatoria* type III effector protein XopJ inhibits protein secretion: evidence for interference with cell wall-associated defense responses. Mol Plant Microbe Interact 22, 655–664.

Bentham, A., Burdett, H., Anderson, P.A., Williams, S.J., and Kobe, B. (2017). Animal NLRs provide structural insights into plant NLR function. Ann Bot 119, 827–702.

Bhandari, D.D., Lapin, D., Kracher, B., von Born, P., Bautor, J., Niefind, K., and Parker, J.E. (2019). An EDS1 heterodimer signalling surface enforces timely reprogramming of immunity genes in Arabidopsis. Nat Commun 10, 772.

Bi, G., Su, M., Li, N., Liang, Y., Dang, S., Xu, J., Hu, M., Wang, J., Zou, M., Deng, Y., Li, Q., Huang, S., Li, J., Chai, J., He, K., Chen, Y.-h., and Zhou, J.-M. (2021). The ZAR1 resistosome is a calcium-permeable channel triggering plant immune signaling. Cell.

Bonardi, V., Tang, S., Stallmann, A., Roberts, M., Cherkis, K., and Dangl, J.L. (2011). Expanded functions for a family of plant intracellular immune receptors beyond specific recognition of pathogen effectors. PNAS 108, 16463–16468.

Burch-Smith, T.M., Schiff, M., Caplan, J.L., Tsao, J., Czymmek, K., and Dinesh-Kumar, S.P. (2007). A novel role for the TIR domain in association with pathogen-derived elicitors. PLoS Biology 5, e68.

Castel, B., Ngou, P.M., Cevik, V., Redkar, A., Kim, D.S., Yang, Y., Ding, P., and Jones, J.D.G. (2018). Diverse NLR immune receptors activate defence via the RPW8-NLR NRG1. New Phytol.

Chen, X., Zuo, S., Schwessinger, B., Chern, M., Canlas, P.E., Ruan, D., Zhou, X., Wang, J., Daudi, A., Petzold, C.J., Heazlewood, J.L., and Ronald, P.C. (2014). An XA21-associated kinase (OsSERK2) regulates immunity mediated by the XA21 and XA3 immune receptors. Mol Plant 7, 874–892.

Chinchilla, D., Zipfel, C., Robatzek, S., Kemmerling, B., Nurnberger, T., Jones, J.D., Felix, G., and Boller, T. (2007). A flagellin-induced complex of the receptor FLS2 and BAK1 initiates plant defence. Nature 448, 497–500.

Collier, S.M., Hamel, L.P., and Moffett, P. (2011). Cell death mediated by the N-terminal domains of a unique and highly conserved class of NB-LRR protein. Mol Plant Microbe Interact 24, 918–931.

Cui, H., Tsuda, K., and Parker, J.E. (2015). Effector-triggered immunity: from pathogen perception to robust defense. Annu Rev Plant Biol 66, 487–511.

Cui, H., Qiu, J., Zhou, Y., Bhandari, D.D., Zhao, C., Bautor, J., and Parker, J.E. (2018). Antagonism of Transcription Factor MYC2 by EDS1/PAD4 Complexes Bolsters Salicylic Acid Defense in Arabidopsis Effector-Triggered Immunity. Mol Plant 11, 1053–1066.

Dongus, J.A., and Parker, J.E. (2021). EDS1 signalling: At the nexus of intracellular and surface receptor immunity. Current Opinion in Plant Biology 62, 102039.

Dongus, J.A., Bhandari, D.D., Patel, M., Archer, L., Dijkgraaf, L., Deslandes, L., Shah, J., and Parker, J.E. (2020). The Arabidopsis PAD4 Lipase-Like Domain Is Sufficient for Resistance to Green Peach Aphid. Molecular Plant-Microbe Interactions 33, 328–335.

Duxbury, Z., Wang, S., MacKenzie, C.I., Tenthorey, J.L., Zhang, X., Huh, S.U., Hu, L., Hill, L., Ngou, P.M., Ding, P., Chen, J., Ma, Y., Guo, H., Castel, B., Moschou, P.N., Bernoux, M., Dodds, P.N., Vance, R.E., and Jones, J.D.G. (2020). Induced proximity of a TIR signaling domain on a plant-mammalian NLR chimera activates defense in plants. PNAS 117, 18832–18839.

Engler, C., Youles, M., Gruetzner, R., Ehnert, T.M., Werner, S., Jones, J.D., Patron, N.J., and Marillonnet, S. (2014). A Golden Gate Modular Cloning Toolbox for Plants. ACS Synthetic Biology.

Essuman, K., Summers, D.W., Sasaki, Y., Mao, X.R., DiAntonio, A., and Milbrandt, J. (2017). The SARM1 Toll/Interleukin-1 Receptor Domain Possesses Intrinsic NAD(+) Cleavage Activity that Promotes Pathological Axonal Degeneration. Neuron 93, 1334-+.

Essuman, K., Summers, D.W., Sasaki, Y., Mao, X., Yim, A.K.Y., DiAntonio, A., and Milbrandt, J. (2018). TIR Domain Proteins Are an Ancient Family of NAD(+)-Consuming Enzymes. Curr Biol 28, 421–430 e424.

Feys, B.J., Wiermer, M., Bhat, R.A., Moisan, L.J., Medina-Escobar, N., Neu, C., Cabral, A., and Parker, J.E. (2005). Arabidopsis SENESCENCE-ASSOCIATED GENE101 stabilizes and signals within an ENHANCED DISEASE SUSCEPTIBILITY1 complex in plant innate immunity. The Plant Cell 17, 2601–2613.

Fradin, E.F., Abd-El-Haliem, A., Masini, L., van den Berg, G.C., Joosten, M.H., and Thomma, B.P. (2011). Interfamily transfer of tomato Ve1 mediates *Verticillium* resistance in Arabidopsis. Plant Physiology 156, 2255–2265.

Fradin, E.F., Zhang, Z., Juarez Ayala, J.C., Castroverde, C.D., Nazar, R.N., Robb, J., Liu, C.M., and Thomma, B.P. (2009). Genetic dissection of Verticillium wilt resistance mediated by tomato Ve1. Plant Physiology 150, 320–332.

Gabriels, S.H., Takken, F.L., Vossen, J.H., de Jong, C.F., Liu, Q., Turk, S.C., Wachowski, L.K., Peters, J., Witsenboer, H.M., de Wit, P.J., and Joosten, M.H. (2006). cDNA-AFLP combined with functional analysis reveals novel genes involved in the hypersensitive response. Mol Plant Microbe Interact 19, 567–576.

Gabriels, S.H.E.J., Vossen, J.H., Ekengren, S.K., van Ooijen, G., Abd-El-Haliem, A.M., van den Berg, G.C.M., Rainey, D.Y., Martin, G.B., Takken, F.L.W., de Wit, P.J.G.M., and Joosten, M.H.A.J. (2007). An NB-LRR protein required for HR signalling mediated by both extra- and intracellular resistance proteins. Plant Journal 50, 14–28.

Gantner, J., Ordon, J., Kretschmer, C., Guerois, R., and Stuttmann, J. (2019). An EDS1-SAG101 Complex Is Essential for TNL-Mediated Immunity in *Nicotiana benthamiana*. The Plant Cell 31, 2456–2474.

Gantner, J., Ordon, J., Ilse, T., Kretschmer, C., Gruetzner, R., Lofke, C., Dagdas, Y., Burstenbinder, K., Marillonnet, S., and Stuttmann, J. (2018). Peripheral infrastructure vectors and an extended set of plant parts for the Modular Cloning system. PLoS ONE 13, e0197185.

Garcia, A.V., Blanvillain-Baufume, S., Huibers, R.P., Wiermer, M., Li, G., Gobbato, E., Rietz, S., and Parker, J.E. (2010). Balanced nuclear and cytoplasmic activities of EDS1 are required for a complete plant innate immune response. PLoS Pathog 6, e1000970.

Goldman, A., Harper, S., and Speicher, D.W. (2016). Detection of Proteins on Blot Membranes. Curr Protoc Protein Sci 86, 10 18 11–10 18 11.

Gomez-Gomez, L., Felix, G., and Boller, T. (1999). A single locus determines sensitivity to bacterial flagellin in *Arabidopsis thaliana*. Plant J 18, 277–284.

Heese, A., Hann, D.R., Gimenez-Ibanez, S., Jones, A.M., He, K., Li, J., Schroeder, J.I., Peck, S.C., and Rathjen, J.P. (2007). The receptor-like kinase SERK3/BAK1 is a central regulator of innate immunity in plants. PNAS 104, 12217–12222.

Hofberger, J.A., Zhou, B., Tang, H., Jones, J.D., and Schranz, M.E. (2014). A novel approach for multi-domain and multi-gene family identification provides insights into evolutionary dynamics of disease resistance genes in core eudicot plants. BMC Genomics 15, 966.

Horsefield, S., Burdett, H., Zhang, X.X., Manik, M.K., Shi, Y., Chen, J., Qi, T.C., Gilley, J., Lai, J.S., Rank, M.X., Casey, L.W., Gu, W.X., Ericsson, D.J., Foley, G., Hughes, R.O., Bosanac, T., von Itzstein, M., Rathjen, J.P., Nanson, J.D., Boden, M., Dry, I.B., Williams, S.J., Staskawicz, B.J., Coleman, M.P., Ve, T., Dodds, P.N., and Kobe, B. (2019). NAD(+) cleavage activity by animal an plant TIR domains in cell death pathways. Science 365.

Hu, G., deHart, A.K., Li, Y., Ustach, C., Handley, V., Navarre, R., Hwang, C.F., Aegerter, B.J., Williamson, V.M., and Baker, B. (2005). *EDS1* in tomato is required for resistance mediated by TIR-class R genes and the receptor-like R gene *Ve*. Plant J 42, 376–391.

Huh, S.U., Cevik, V., Ding, P., Duxbury, Z., Ma, Y., Tomlinson, L., Sarris, P.F., and Jones, J.D.G. (2017). Protein-protein interactions in the RPS4/RRS1 immune receptor complex. PLoS Pathog 13, e1006376.

Jacob, P., Kim, N.H., Wu, F., El-Kasmi, F., Chi, Y., Walton, W.G., Furzer, O.J., Lietzan, A.D., Sunil, S., Kempthorn, K., Redinbo, M.R., Pei, Z.M., Wan, L., and Dangl, J.L. (2021). Plant “helper” immune receptors are Ca(2+)-permeable nonselective cation channels. Science 373, 420–425.

Johanndrees, O., Baggs, E.L., Uhlmann, C., Locci, F., Läßle, H.L., Melkonian, K., Käufer, K., Dongus, J.A., Nakagami, H., Krasileva, K.V., Parker, J.E., and Lapin, D. (2021). Differential *EDS1* requirement for cell death activities of plant TIR-domain proteins. bioRxiv, 2021.2011.2029.470438.

Kourelis, J., Contreras, M.P., Harant, A., Adachi, H., Derevnina, L., Wu, C.-H., and Kamoun, S. (2021). The helper NLR immune protein NRC3 mediates the hypersensitive cell death caused by the cell-surface receptor Cf-4. bioRxiv, 2021.2009.2028.461843.

Landeo Villanueva, S., Malvestiti, M.C., van Ieperen, W., Joosten, M., and van Kan, J.A.L. (2021). Red light imaging for programmed cell death visualization and quantification in plant-pathogen interactions. Mol Plant Pathol 22, 361–372.

Lapin, D., Bhandari, D.D., and Parker, J.E. (2020). Origins and Immunity Networking Functions of EDS1 Family Proteins. Annu Rev Phytopathol 58, 253–276.

Lapin, D., Kovacova, V., Sun, X., Dongus, J.A., Bhandari, D.D., von Born, P., Bautor, J., Guarneri, N., Rzemieniewski, J., Stuttmann, J., Beyer, A., and Parker, J.E. (2019). A coevolved EDS1-SAG101-NRG1 module mediates cell death signaling by TIR-domain immune receptors. The Plant Cell.

Lewis, J.D., Abada, W., Ma, W., Guttman, D.S., and Desveaux, D. (2008). The HopZ family of *Pseudomonas syringae* type III effectors require myristoylation for virulence and avirulence functions in *Arabidopsis thaliana*. J Bacteriol 190, 2880–2891.

Liebrand, T.W., van den Berg, G.C., Zhang, Z., Smit, P., Cordewener, J.H., America, A.H., Sklenar, J., Jones, A.M., Tameling, W.I., Robatzek, S., Thomma, B.P., and Joosten, M.H. (2013). Receptor-like kinase SOBIR1/EVR interacts with receptor-like proteins in plant immunity against fungal infection. PNAS 110, 10010–10015.

Liu, Y., Zeng, Z., Zhang, Y.-M., Li, Q., Jiang, X.-M., Jiang, Z., Tang, J.-H., Chen, D., Wang, Q., Chen, J.-Q., and Shao, Z.-Q. (2021). An angiosperm *NLR* Atlas reveals that *NLR* gene reduction is associated with ecological specialization and signal transduction component deletion. Molecular Plant.

Logemann, E., Birkenbihl, R.P., Ulker, B., and Somssich, I.E. (2006). An improved method for preparing Agrobacterium cells that simplifies the Arabidopsis transformation protocol. Plant Methods 2, 16.

Louis, J., Gobbato, E., Mondal, H.A., Feys, B.J., Parker, J.E., and Shah, J. (2012). Discrimination of Arabidopsis PAD4 activities in defense against green peach aphid and pathogens. Plant Physiology 158, 1860–1872.

Lu, Y., and Tsuda, K. (2021). Intimate Association of PRR- and NLR-Mediated Signaling in Plant Immunity. Mol Plant Microbe Interact 34, 3–14.

Ma, S., Lapin, D., Liu, L., Sun, Y., Song, W., Zhang, X., Logemann, E., Yu, D., Wang, J., Jirschitzka, J., Han, Z., Schulze-Lefert, P., Parker, J.E., and Chai, J. (2020). Direct pathogen-induced assembly of an NLR immune receptor complex to form a holoenzyme. Science 370.

Mahdi, L.K., Huang, M., Zhang, X., Nakano, R.T., Kopp, L.B., Saur, I.M.L., Jacob, F., Kovacova, V., Lapin, D., Parker, J.E., Murphy, J.M., Hofmann, K., Schulze-Lefert, P., Chai, J., and Maekawa, T. (2020). Discovery of a Family of Mixed Lineage Kinase Domain-like Proteins in Plants and Their Role in Innate Immune Signaling. Cell Host & Microbe 28, 813–824.e816.

Martin, R., Qi, T., Zhang, H., Liu, F., King, M., Toth, C., Nogales, E., and Staskawicz, B.J. (2020). Structure of the activated ROQ1 resistosome directly recognizing the pathogen effector XopQ. Science 370, eabd9993.

Mindrebo, J.T., Nartey, C.M., Seto, Y., Burkartl, M.D., and Noel, J.P. (2016). Unveiling the functional diversity of the alpha/beta hydrolase superfamily in the plant kingdom. Curr Opin Struc Biol 41, 256–257.

Narusaka, M., Shirasu, K., Noutoshi, Y., Kubo, Y., Shiraishi, T., Iwabuchi, M., and Narusaka, Y. (2009). RRS1 and RPS4 provide a dual Resistance-gene system against fungal and bacterial pathogens. The Plant Journal 60, 218–226.

Ngou, B.P.M., Ahn, H.K., Ding, P., and Jones, J.D.G. (2021). Mutual potentiation of plant immunity by cell-surface and intracellular receptors. Nature 592, 110–115.

Ofir, G., Herbst, E., Baroz, M., Cohen, D., Millman, A., Doron, S., Tal, N., Malheiro, D.B.A., Malitsky, S., Amitai, G., and Sorek, R. (2021). Antiviral activity of bacterial TIR domains via immune signalling molecules. Nature 600, 116–120.

Ordon, J., Martin, P., Erickson, J.L., Ferik, F., Balcke, G., Bonas, U., and Stuttmann, J. (2021). Disentangling cause and consequence: genetic dissection of the *DANGEROUS MIX2* risk locus, and activation of the DM2h NLR in autoimmunity. Plant Journal.

Ordon, J., Gantner, J., Kemna, J., Schwalgun, L., Reschke, M., Streubel, J., Boch, J., and Stuttmann, J. (2017). Generation of chromosomal deletions in dicotyledonous plants employing a user-friendly genome editing toolkit. Plant Journal 89, 155–168.

Postma, J., Liebrand, T.W., Bi, G., Evrard, A., Bye, R.R., Mbengue, M., Kuhn, H., Joosten, M.H., and Robatzek, S. (2016). Avr4 promotes Cf-4 receptor-like protein association with the BAK1/SERK3 receptor-like kinase to initiate receptor endocytosis and plant immunity. New Phytol 210, 627–642.

Prautsch, J., Erickson, J.L., Özyürek, S., Gormanns, R., Franke, L., Parker, J.E., Stuttmann, J., and Schattat, M.H. (2021). XopQ induced stromule formation in *Nicotiana benthamiana* is causally linked to ETI signaling and depends on ADR1 and NRG1. bioRxiv, 2021.2012.2006.471425.

Pruitt, R.N., Locci, F., Wanke, F., Zhang, L., Saile, S.C., Joe, A., Karelina, D., Hua, C., Frohlich, K., Wan, W.L., Hu, M., Rao, S., Stolze, S.C., Harzen, A., Gust, A.A., Harter, K., Joosten, M., Thomma, B., Zhou, J.M., Dangl, J.L., Weigel, D., Nakagami, H., Oecking, C., Kasmi, F.E., Parker, J.E., and Nurnberger, T. (2021). The EDS1-PAD4-ADR1 node mediates Arabidopsis pattern-triggered immunity. Nature.

Qi, T., Seong, K., Thomazella, D.P.T., Kim, J.R., Pham, J., Seo, E., Cho, M.J., Schultink, A., and Staskawicz, B.J. (2018). NRG1 functions downstream of EDS1 to regulate TIR-NLR-mediated plant immunity in Nicotiana benthamiana. PNAS.

Rauwerdink, A., and Kazlauskas, R.J. (2015). How the Same Core Catalytic Machinery Catalyzes 17 Different Reactions: the Serine-Histidine-Aspartate Catalytic Triad of α/β-Hydrolase Fold Enzymes. ACS Catalysis 5, 6153–6176.

Saijo, Y., Loo, E.P., and Yasuda, S. (2018). Pattern recognition receptors and signaling in plant-microbe interactions. Plant Journal 93, 592–613.

Saile, S.C., Jacob, P., Castel, B., Jubic, L.M., Salas-Gonzales, I., Backer, M., Jones, J.D.G., Dangl, J.L., and El Kasmi, F. (2020). Two unequally redundant “helper” immune receptor families mediate *Arabidopsis thaliana* intracellular “sensor” immune receptor functions. PLoS Biology 18, e3000783.

Saile, S.C., Ackermann, F.M., Sunil, S., Keicher, J., Bayless, A., Bonardi, V., Wan, L., Doumane, M., Stobbe, E., Jaillais, Y., Caillaud, M.C., Dangl, J.L., Nishimura, M.T., Oecking, C., and El Kasmi, F. (2021). Arabidopsis ADR1 helper NLR immune receptors localize and function at the plasma membrane in a phospholipid dependent manner. New Phytol.

Saucet, S.B., Ma, Y., Sarris, P.F., Furzer, O.J., Sohn, K.H., and Jones, J.D. (2015). Two linked pairs of Arabidopsis TNL resistance genes independently confer recognition of bacterial effector AvrRps4. Nat Commun 6, 6338.

Schultink, A., Qi, T., Lee, A., Steinbrenner, A.D., and Staskawicz, B. (2017). Roq1 mediates recognition of the *Xanthomonas* and *Pseudomonas* effector proteins XopQ and HopQ1. Plant Journal.

Seto, Y., Yasui, R., Kameoka, H., Tamiru, M., Cao, M., Terauchi, R., Sakurada, A., Hirano, R., Kisugi, T., Hanada, A., Umehara, M., Seo, E., Akiyama, K., Burke, J., Takeda-Kamiya, N., Li, W., Hirano, Y., Hakoshima, T., Mashiguchi, K., Noel, J.P., Kyozuka, J., and Yamaguchi, S. (2019). Strigolactone perception and deactivation by a hydrolase receptor DWARF14. Nat Commun 10, 191.

Shimada, A., Ueguchi-Tanaka, M., Nakatsu, T., Nakajima, M., Naoe, Y., Ohmiya, H., Kato, H., and Matsuoka, M. (2008). Structural basis for gibberellin recognition by its receptor GID1. Nature 456, 520–523.

Shimada, T.L., Shimada, T., and Hara-Nishimura, I. (2010). A rapid and non-destructive screenable marker, FAST, for identifying transformed seeds of Arabidopsis thaliana. Plant Journal 61, 519–528.

Sinapidou, E., Williams, K., Nott, L., Bahkt, S., Tor, M., Crute, I., Bittner-Eddy, P., and Beynon, J. (2004). Two TIR:NB:LRR genes are required to specify resistance to *Peronospora parasitica* isolate Cala2 in Arabidopsis. Plant J 38, 898–909.

Stuttmann, J., Hubberten, H.M., Rietz, S., Kaur, J., Muskett, P., Guerois, R., Bednarek, P., Hoefgen, R., and Parker, J.E. (2011). Perturbation of Arabidopsis amino acid metabolism causes incompatibility with the adapted biotrophic pathogen *Hyaloperonospora arabidopsidis*. The Plant Cell 23, 2788–2803.

Stuttmann, J., Peine, N., Garcia, A.V., Wagner, C., Choudhury, S.R., Wang, Y., James, G.V., Griebel, T., Alcazar, R., Tsuda, K., Schneeberger, K., and Parker, J.E. (2016). *Arabidopsis thaliana DM2h* (*R8*) within the Landsberg *RPP1-like* Resistance Locus Underlies Three Different Cases of EDS1-Conditioned Autoimmunity. PLoS Genet 12, e1005990.

Stuttmann, J., Barthel, K., Martin, P., Ordon, J., Erickson, J.L., Herr, R., Ferik, F., Kretschmer, C., Berner, T., Keilwagen, J., Marillonnet, S., and Bonas, U. (2021). Highly efficient multiplex editing: one-shot generation of 8x *Nicotiana benthamiana* and 12x Arabidopsis mutants. Plant Journal 106, 8–22.

Sun, X., Lapin, D., Feehan, J.M., Stolze, S.C., Kramer, K., Dongus, J.A., Rzemieniewski, J., Blanvillain-Baufumé, S., Harzen, A., Bautor, J., Derbyshire, P., Menke, F.L.H., Finkemeier, I., Nakagami, H., Jones, J.D.G., and Parker, J.E. (2021). Pathogen effector recognition-dependent association of NRG1 with EDS1 and SAG101 in TNL receptor immunity. Nature Communications 12, 3335.

Takemoto, D., Rafiqi, M., Hurley, U., Lawrence, G.J., Bernoux, M., Hardham, A.R., Ellis, J.G., Dodds, P.N., and Jones, D.A. (2012). N-terminal motifs in some plant disease resistance proteins function in membrane attachment and contribute to disease resistance. Mol Plant Microbe Interact 25, 379–392.

Tao, Y., Xie, Z.Y., Chen, W.Q., Glazebrook, J., Chang, H.S., Han, B., Zhu, T., Zou, G.Z., and Katagiri, F. (2003). Quantitative nature of Arabidopsis responses during compatible and incompatible interactions with the bacterial pathogen *Pseudomonas syringae*. The Plant Cell 15, 317–330.

Thieme, F., Koebnik, R., Bekel, T., Berger, C., Boch, J., Buttner, D., Caldana, C., Gaigalat, L., Goesmann, A., Kay, S., Kirchner, O., Lanz, C., Linke, B., McHardy, A.C., Meyer, F., Mittenhuber, G., Nies, D.H., Niesbach-Klosgen, U., Patschkowski, T., Ruckert, C., Rupp, O., Schneiker, S., Schuster, S.C., Vorholter, F.J., Weber, E., Puhler, A., Bonas, U., Bartels, D., and Kaiser, O. (2005). Insights into genome plasticity and pathogenicity of the plant pathogenic bacterium *Xanthomonas campestris* pv. *vesicatoria* revealed by the complete genome sequence. J Bacteriol 187, 7254–7266.

Thomma, B.P.H.J., Nürnberger, T., and Joosten, M.H.A.J. (2011). Of PAMPs and effectors: the blurred PTI-ETI dichotomy. The Plant Cell 23, 4–15.

Tian, H., Wu, Z., Chen, S., Ao, K., Huang, W., Yaghmaiean, H., Sun, T., Xu, F., Zhang, Y., Wang, S., Li, X., and Zhang, Y. (2021). Activation of TIR signalling boosts pattern-triggered immunity. Nature.

Van de Weyer, A.L., Monteiro, F., Furzer, O.J., Nishimura, M.T., Cevik, V., Witek, K., Jones, J.D.G., Dangl, J.L., Weigel, D., and Bemm, F. (2019). A Species-Wide Inventory of NLR Genes and Alleles in *Arabidopsis thaliana*. Cell 178, 1260–1272 e1214.

Van der Hoorn, R.A., Laurent, F., Roth, R., and De Wit, P.J. (2000). Agroinfiltration is a versatile tool that facilitates comparative analyses of Avr9/Cf-9-induced and Avr4/Cf-4-induced necrosis. Mol Plant Microbe Interact 13, 439–446.

Voss, M., Toelzer, C., Bhandari, D.D., Parker, J.E., and Niefind, K. (2019). Arabidopsis immunity regulator EDS1 in a PAD4/SAG101-unbound form is a monomer with an inherently inactive conformation. J Struct Biol 208, 107390.

Wagner, S., Stuttmann, J., Rietz, S., Guerois, R., Brunstein, E., Bautor, J., Niefind, K., and Parker, J.E. (2013). Structural basis for signaling by exclusive EDS1 heteromeric complexes with SAG101 or PAD4 in plant innate immunity. Cell Host Microbe 14, 619–630.

Wan, L., Essuman, K., Anderson, R.G., Sasaki, Y., Monteiro, F., Chung, E.H., Nishimura, E.O., DiAntonio, A., Milbrandt, J., Dangl, J.L., and Nishimura, M.T. (2019). TIR domains of plant immune receptors are NAD(+)-cleaving enzymes that promote cell death. Science 365.

Wang, J., Hu, M., Wang, J., Qi, J., Han, Z., Wang, G., Qi, Y., Wang, H.W., Zhou, J.M., and Chai, J. (2019). Reconstitution and structure of a plant NLR resistosome conferring immunity. Science 364.

Wang, Y., Xu, Y., Sun, Y., Wang, H., Qi, J., Wan, B., Ye, W., Lin, Y., Shao, Y., Dong, S., Tyler, B.M., and Wang, Y. (2018). Leucine-rich repeat receptor-like gene screen reveals that *Nicotiana* RXEG1 regulates glycoside hydrolase 12 MAMP detection. Nature Communications 9, 594.

Wen, W., Meinkoth, J.L., Tsien, R.Y., and Taylor, S.S. (1995). Identification of a signal for rapid export of proteins from the nucleus. Cell 82, 463–473.

Wirthmueller, L., Zhang, Y., Jones, J.D., and Parker, J.E. (2007). Nuclear accumulation of the Arabidopsis immune receptor RPS4 is necessary for triggering EDS1-dependent defense. Curr Biol 17, 2023–2029.

Wu, C.-H., Abd-El-Haliem, A., Bozkurt, T.O., Belhaj, K., Terauchi, R., Vossen, J.H., and Kamoun, S. (2017). NLR network mediates immunity to diverse plant pathogens. PNAS 114, 8113–8118.

Wu, Z., Tian, L., Liu, X., Zhang, Y., and Li, X. (2021). TIR signal promotes interactions between lipase-like proteins and ADR1-L1 receptor and ADR1-L1 oligomerization. Plant Physiology 187, 681–686.

Wu, Z., Li, M., Dong, O.X., Xia, S., Liang, W., Bao, Y., Wasteneys, G., and Li, X. (2018). Differential regulation of TNL-mediated immune signaling by redundant helper CNLs. New Phytol.

Xiao, S., Calis, O., Patrick, E., Zhang, G., Charoenwattana, P., Muskett, P., Parker, J.E., and Turner, J.G. (2005). The atypical resistance gene, *RPW8*, recruits components of basal defence for powdery mildew resistance in Arabidopsis. Plant Journal. 42, 95–110.

Yu, D., Song, W., Tan, E.Y.J., Liu, L., Cao, Y., Jirschitzka, J., Li, E., Logemann, E., Xu, C., Huang, S., Jia, A., Chang, X., Han, Z., Wu, B., Schulze-Lefert, P., and Chai, J. (2021). TIR domains of plant immune receptors are 2′,3′-cAMP/cGMP synthetases mediating cell death. bioRxiv, 2021.2011.2009.467869.

Yuan, M., Jiang, Z., Bi, G., Nomura, K., Liu, M., Wang, Y., Cai, B., Zhou, J.M., He, S.Y., and Xin, X.F. (2021). Pattern-recognition receptors are required for NLR-mediated plant immunity. Nature 592, 105–109.

Zipfel, C. (2014). Plant pattern-recognition receptors. Trends Immunol 35, 345–351.

